# Crosslinker Structure Modulates Bulk Mechanical Properties and Dictates hMSC Behavior on Hyaluronic Acid Hydrogels

**DOI:** 10.1101/2022.08.03.502671

**Authors:** Logan D. Morton, David A. Castilla-Casadiego, Ajay C. Palmer, Adrianne M. Rosales

**Author notes:** Corresponding author: Adrianne M. Rosales, McKetta Department of Chemical Engineering, University of Texas at Austin, Austin, TX, USA.

## Abstract

Synthetic hydrogels are attractive platforms due in part to their highly tunable mechanics, which impact cell behavior and secretory profile. These mechanics are often controlled by altering the number of crosslinks or the total polymer concentration in the gel, leading to structure-property relationships that inherently couple network connectivity to the overall modulus. In contrast, the native extracellular matrix (ECM) contains structured biopolymers that enable stiff gels even at low polymer content, facilitating 3D cell culture and permeability of soluble factors. To mimic the hierarchical order of natural ECM, this work describes a synthetic hydrogel system in which mechanics are tuned using the structure of sequence-defined peptoid crosslinkers, while fixing network connectivity. Peptoid crosslinkers with different secondary structures are investigated: 1) a helical, molecularly stiff peptoid, 2) a non-helical, less stiff peptoid, and 3) an unstructured, relatively flexible peptoid. Bulk hydrogel storage modulus increases when crosslinkers of higher chain stiffness are used. In-vitro studies assess the viability, proliferation, cell morphology, and immunomodulatory activity of human mesenchymal stem cells (hMSCs) on each hydrogel substrate. Matrix mechanics regulate the morphology of hMSCs on the developed substrates, and all of the hydrogels studied upregulate IDO production over culture on TCP. Softer substrates further this upregulation to a plateau. Overall, this system offers a biomimetic strategy for decoupling hydrogel storage modulus from network connectivity, enabling systematic study of biomaterial properties on hMSC behavior and enhancement of cellular functionality for therapeutic applications.

**Graphical Abstract:** 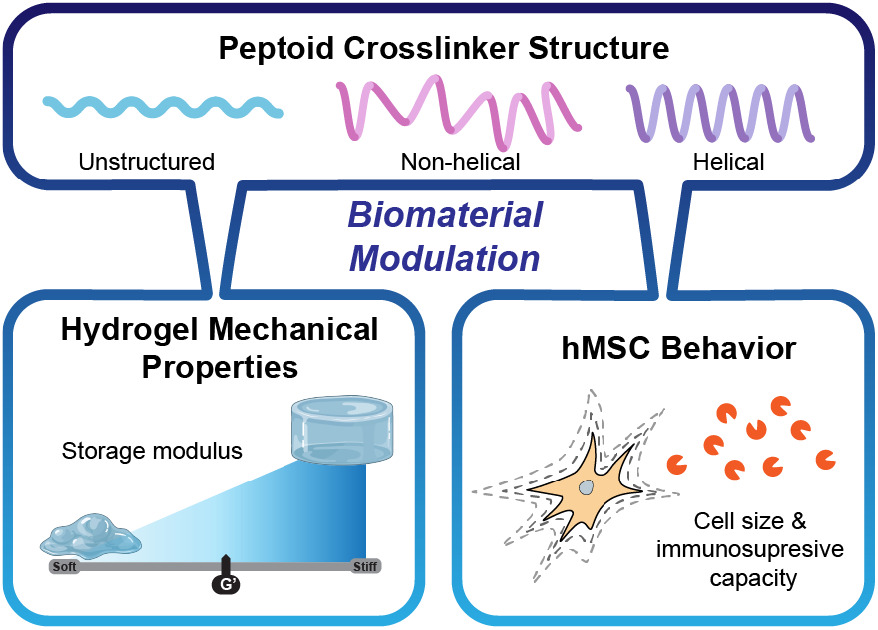

## Introduction

Synthetic hydrogels have garnered intense interest as extracellular matrix (ECM) mimics, yet still do not fully replicate the hierarchical nature of biological tissue, which limits their biological efficacy as stem cell culture scaffolds for tissue engineering and regenerative medicine. The native ECM is made up of hierarchical biopolymers with precise sequences and chain structures that yield high molecular rigidity.^[1–4]^ These structures result in stiff microenvironments at relatively low polymer concentrations. Synthetic hydrogels, meanwhile, typically utilize changes in polymer concentration or crosslinking stoichiometry to modulate mechanics with flexible, disordered polymers.^[5]^ While this approach yields tunable and predictable moduli, it inherently links mechanics to other parameters of the hydrogel system, such as network connectivity and mesh size, and therefore affects permeability to cytokines and growth factors.^[6–10]^ This inherent coupling may diminish the efficacy of synthetic hydrogels as stem cell culture scaffolds by hindering the concentration or availability of cell signaling factors, as well as disrupting the matrix remodeling processes.

To decouple mechanics from network connectivity, recent efforts have investigated the incorporation of stiff, biomimetic chains into synthetic hydrogel formulations.^[11–15]^ Helical polypeptides were incorporated into poly(ethylene glycol, PEG)-based hydrogels as crosslinkers, yielding higher elastic moduli than hydrogels with non-helical polypeptide crosslinkers due to the increased persistence length of the helices.^[11]^ In these studies, hydrogels formed with helical crosslinkers facilitated attachment of human mesenchymal stromal cells (hMSCs)^[13]^ and endothelial cells^[14]^ by increasing local stiffness at the nanoscale. In a complementary strategy, collagen mimetic peptides were incorporated into synthetic PEG hydrogels, capturing the multiscale structure of collagen-rich tissue and leading to cell morphologies that resembled those in traditional collagen platforms.^[15]^ While both of the aforementioned systems leveraged biological motifs for stiff secondary structures, it is important to note that they are sensitive to environmental and processing conditions that affect hydrogen bonding interactions. In a non-biological system, it was shown that controlling the stereochemistry of alkene linkages in a PEG hydrogel resulted in control over bulk mechanics, with larger *cis* contents leading to lower elastic moduli.^[12]^ Again, control over stereochemistry was achieved by adjusting solvent polarity and basicity during hydrogel fabrication. To avoid this dependence, more robust synthetic strategies to control chain rigidity and structure are needed.

Peptoids are a class of non-natural molecules capable of forming biomimetic secondary structures based on their sequence definition, which arises from primary amine submonomers.^[16– 18]^ These secondary structures include helices,^[19–23]^ which have been shown to have higher persistence lengths^[24]^ and shorter end-to-end distances^[19,25]^ than non-helical analogs. Because peptoids form polyproline type I helices through sterics,^[21,26]^ they are less sensitive to temperature and solvent conditions than peptide helices, which rely on hydrogen bonding interactions.^[27]^ Previous work has shown helical peptoids can alter the self-assembled nanostructure of block copolymers,^[28,29]^ and that they have more local chain stiffness and compact chain conformations compared to racemic analogs.^[25,30]^ In addition, we previously showed that increasing the length of helical peptoid crosslinkers increased the elastic moduli of PEG hydrogels, breaking the trend predicted by rubber elasticity theory for flexible polymer networks.^[31]^ Thus, peptoids are an excellent model system for probing the effect of crosslinker chain structure on bulk mechanics for engineered ECM applications.

In this work, we investigated the impact of peptoid crosslinker secondary structure on hydrogel mechanics and whether the resulting changes in mechanics affected attached cell behavior. Our strategy was to decouple hydrogel storage modulus (*G’*) from network connectivity using a suite of sequence defined peptoids with different chain structures: a helical sequence, a non-helical but chemically similar sequence, and an unstructured sequence. This series of peptoids enabled a highly tunable crosslinker stiffness, expanding the toolkit of available molecules for synthetic ECM. To determine the applicability of this system for modulating cell-ECM interactions, we seeded human mesenchymal stromal cells (hMSCs) on the hydrogels, since these cells are promising for allogenic cell therapies and exhibit immunomodulatory functions that are sensitive to matrix mechanics.^[32–36]^ We found that the range of stiffness was sufficient to affect cell morphology, proliferation, and immunomodulatory activity, with softer hydrogels producing more therapeutically potent cells. Altogether, this study highlights a unique way to regulate bulk mechanics using a property that is often overlooked or unachievable in synthetic ECM: molecular rigidity.

## Materials and Methods

### Peptoid Synthesis

The helical and non-helical peptoids were synthesized using Rink Amide polystyrene resin (0.43-0.49 mmol/g from Chem-Impex International, Inc.) on a Prelude X automated peptide synthesizer (Gyros Protein Technologies) at a scale of 250 μM according to previously published submonomer methods.^[16,37]^ For each synthesis, fresh bromoacetylation reagent was made by dissolving 1.2 M bromoacetic acid (>99%, Fisher Scientific) in dimethylformamide (DMF) (99.8%, Fisher Scientific). All displacement steps utilized 2 M primary amines (Trt-Cysteamine*HCl (>99%), (S)-(-)-1-Phenylethylamine (>99%), L-Alanine-tert butyl ester*HCl (>97.5%) (Chem-Impex International, Inc.) and α-methylbenzylamine (99%, Sigma Aldrich)) in N-methyl-2-pyrrolidone (NMP) (>99%, Fisher Scientific), after freebasing when necessary.

The unstructured peptoid was synthesized using Rink Amide polystyrene resin (0.49 mmol/g from Chem-Impex International, Inc.) on a Prelude X automated peptide synthesizer (Gyros Protein Technologies). Traditional Fmoc-mediated coupling methods were used at five-fold molar excess of Fmoc-*S*-trityl-L-Cysteine (99.44%) and Fmoc-*N*-methylglycine (99.51%) (Chem-Impex International, Inc.), and O-(1H-6-Chlorobenzotriazole-1-yl)-1,1,3,3-tetramethyluronium hexafluorophosphate (HCTU) coupling reagent (99.9%, Chem-Impex International, Inc.), and a ten-fold excess of N-methylmorpholine (99%, Sigma Aldrich). Coupling steps were performed twice.

Once complete, all syntheses were cleaved from resin with 5 mL of 92.5:2.5:2.5:2.5 trifluoroacetic acid (>95%, Fisher Scientific):DI water:triisopropylsilane (98%, Fisher Scientific):1,2-ethanedithiol (>98%, Sigma Aldrich) for 4 hours. Prior to purification, the peptoids were dissolved at 10 mg/mL in 50:50 acetonitrile (>99.9%, Fisher Scientific):DI water with 0.1% trifluoroacetic acid. The crude peptoids were purified with a C18 column on a Dionex UltiMate 3000 UHPLC using a 50 min gradient of acetonitrile in water (50% - 100%, 10 mL/min). After lyophilization, masses were verified via matrix-assisted laser desorption/ionization-time of flight (MALDI-TOF) mass spectrometry (Applied Biosystems - Voyager-DE™ PRO) (**Figure S1**) and purity was confirmed via analytical HPLC (**Figure S2**).

### Peptide Synthesis

Peptides were synthesized using Rink Amide polystyrene resin (0.43-0.49 mmol/g, Chem-Impex International, Inc.) on a Prelude X automated peptide synthesizer (Gyros Protein Technologies). Traditional Fmoc-mediated coupling methods were used at five-fold molar excess of amino acids and O-(1H-6-Chlorobenzotriazole-1-yl)-1,1,3,3-tetramethyluronium hexafluorophosphate (HCTU) coupling reagent, and a ten-fold excess of N-methylmorpholine. Coupling steps were performed twice. Once complete, both syntheses were cleaved from resin with 10 mL of 95:2.5:2.5 trifluoroacetic acid:water:triisopropylsilane for 4 hours. Prior to purification, the peptides were dissolved at 10 mg/mL in 20:80 acetonitrile:water with 0.1% trifluoroacetic acid. The crude peptides were purified with a C18 column on a Dionex UltiMate 3000 UHPLC using a 40 min gradient of acetonitrile in water (20% - 100%, 10 mL/min). After lyophilization, masses were verified via MALDI-TOF (**Figure S3**). The sequences of the peptides synthesized are KCGGIQQWGPCK (for the peptide crosslinker control) and GCGYGRGDSPG (for cell adhesion).

### Synthesis of Norbornene-functionalized Hyaluronic Acid (NorHA)

NorHA was synthesized following the procedure from *Wade et al*.^[38]^ with small modifications. Briefly, hyaluronic acid (HA) was first converted to a tetrabutylammonium salt (HA-TBA). Sodium HA (Na-HA, 72 kDa, Lifecore) was dissolved in DI water at 2 wt% with Dowex 50W ion exchange resin (Sigma Aldrich) (5:1 resin to HA by weight) and stirred for 5 hours. The resin was filtered out, and the solution was titrated to pH 7.05 with tetrabutylammonium hydroxide, frozen, and lyophilized. Next, HA-TBA (2 wt%), 5-norbornene-2-carboxylic acid (99%, Fisher Scientific) (12:1 to HA-TBA repeat unit), and 4-(dimethylamino)pyridine (Fisher Scientific) (3:1 to HA-TBA repeat unit) were dissolved in anhydrous dimethyl sulfoxide (DMSO) (>99%, Fisher Scientific) under argon at 45°C. Next, di-tert-butyl dicarbonate (Boc_2_O, 99%, Fisher Scientific) (1.6:1 to HA-TBA repeat unit) was added to the flask via syringe, and the reaction proceeded overnight. The reaction was then quenched with 5x excess DI water and dialyzed for three days. On day 3, 1 g NaCl per 100 mL was added, and the solution was precipitated into cold (4°C) acetone (>99%, Fisher Scientific). The precipitate was spun down and re-dissolved into DI water, dialyzed for an additional week, frozen and lyophilized. Proton nuclear magnetic resonance spectroscopy (^1^H NMR) confirmed ∼32% of HA repeat units had been functionalized with norbornene (**Figure S4**).

### Hydrogel Preparation

All gelation solutions were 50% DMSO and 50% phosphate buffered saline with 0.05 wt% Lithium phenyl-2,4,6-trimethylbenzoylphosphinate (LAP). All hydrogels were 3 wt% NorHA crosslinked at 1:1 thiol:ene ratio with either the peptide or peptoid crosslinker. Gelation was induced via exposure to 365 nm light (10 mW/cm^2^, 3 min). For experiments with live cells, the hydrogels were functionalized with 2 mM of the cell adhesive peptide to allow for attachment, and hydrogels were washed several times and swollen overnight in Dulbeccco’s phosphate-buffered saline (DPBS without Ca^++^ and Mg^++^, Corning, Cat. #21-031-CV) to remove the remaining DMSO prior to cell seeding. The hydrogels were then incubated for 1 hour in media before seeding the hMSCs.

### Circular Dichroism (CD)

CD spectra were acquired from 180 to 280 nm using a Jasco J-815 CD Spectropolarimeter at 25°C. Samples (200 μL) were prepared at 2 mM in 20% acetonitrile in water for each of the peptoids. A spectrum of pure 20% acetonitrile in water was subtracted from each sample spectrum to remove any baseline. A quartz cuvette with a 1 mm path length was used for all samples. Per residue molar ellipticity was calculated using the following equation:

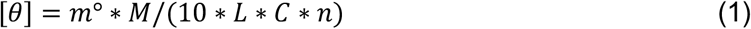

where *m*° is the CD given in millidegrees, *M* is the molecular weight (g/mol), *L* is the path length of the cell in cm, *C* is the concentration of peptoid in g/L, and *n* is the number of monomers.

### Rheometry

A Discovery HR-2 rheometer (TA Instruments) was used to perform all *in situ* gelation mechanics measurements. An 8 mm flat stainless-steel geometry and a UV-transparent quartz plate were utilized for all experiments. Briefly, the macromer solution (15 µL) was pipetted onto the quartz plate, which was connected via liquid filled light guide to a mercury lamp (Omnicure Series 1500) fitted with a 365 nm filter. The macromer solution was exposed *in situ* to 365 nm light (10 mW/cm^2^, 3 min) to cause gelation. Time sweeps were performed at 1 rad/s and 1% strain.

### Nanoindentation

Hydrogels for nanoindentation were fabricated on coverslips functionalized with 3-(Trimethoxysilyl)propyl methacrylate (Fisher Scientific) as previously described.^[39]^ Briefly, coverslips were immersed in 10 M NaOH for 20 minutes in a petri dish, thoroughly rinsed with DI water, then immersed in a solution of 0.4% 3-(Trimethoxysilyl)propyl methacrylate in acetone for 20 minutes. A silicon spacer was placed on top of a hydrophobic slide and biopunched to create the geometry of the hydrogel. The macromer solution (25 µL) was pipetted directly onto the hydrophobic slide and formed into a cylindrical disk 8 mm in diameter. The coverslip was then placed on top and pressed firmly to contact the solution. Gelation was induced via exposure to 365 nm light (10 mW/cm^2^, 3 min) and the hydrophobic slide was removed by submerging under water. The hydrogels were then allowed to swell for 3 days. All samples were measured submerged in PBS on a Piuma Nanoindentor, and elastic moduli were taken directly from the Hertzian fit provided from the software.

### Cell culture

hMSCs (RoosterBio, Cat. #MSC-031, Tissue origin: Human Bone Marrow) were used between passages 3−5 for all the experiments. hMSCs from a healthy 29-year-old male (Donor 1; Lot. 310277) and 23-year-old male (Donor 2; Lot. 00257) were used. The product specification sheet provided by the vendor shows that these cells conserved adipogenic and osteogenic potential, and were positive for CD90 and CD166 surface markers, and negative for CD34 and CD45, as tested by multiplex flow cytometry. hMSCs were grown in alpha-minimum essential media (1×) (supplemented with L-glutamine, ribonucleosides, and deoxyribonucleosides) (Gibco, Cat. #12561-056) containing 20% fetal bovine serum (Gibco, Cat. #12662029), 1.2% penicillin-streptomycin (Corning, Cat. #30002CI), and 1.2% L-glutamine (Corning, Cat. #25005CI). For some in-vitro studies as noted, the medium was supplemented with Human Interferon-gamma Recombinant Protein (IFN-γ) (ThermoFisher, Cat. #. PHC4031) at a concentration of 50 ng/mL. Conditions with medium supplemented with and without IFN-γ were designated as (+) IFN-γ and (−) IFN-γ, respectively.

### Cell viability

An MTT assay (InvitrogenTM, Cat. #V13154) was used to measure cellular metabolic activity as an indicator of cell viability for hMSCs (3,000 cells/cm^2^) seeded on the peptoid-crosslinked hydrogels with and without IFN-γ supplementation. A peptide-crosslinked hydrogel was used as a control. All the conditions were assessed in the presence and absence of IFN-γ at a concentration of 50 ng/mL. Cell viability was measured after 3 days of culture at 37°C and 5% CO_2_, as described in our previous work.^[31]^ Briefly, the cell culture medium was removed, and 100 µL of serum-free media and 10 µL of 3-(4,5-dimethylthiazol-2-yl)-2,5-diphenyltetrazolium bromide (MTT) were added to each well and allowed to incubate for 4 hours. Sodium dodecyl sulfate (SDS) was subsequently added and incubated for another 18 hours to lyse the cells and solubilize the metabolized MTT. The absorbance from all wells was measured using a Synergy H1 Hybrid Multi-Mode Reader at a wavelength of 570 nm.

### Cell Proliferation

A Click-iT Plus EdU Alexa Fluor 488 Imaging Kit from Invitrogen (cat. no. C10337) was used to measure the ability of cells to proliferate. hMSCs (10000 cells/cm^2^) were seeded on each hydrogel condition. A peptide-crosslinked hydrogel and tissue culture plastic (TCP) were used as controls. All the conditions were assessed in the presence and absence of IFN-γ at a concentration of 50 ng/mL. After three days of culture, half of the medium was removed, and EdU solution was added according to the protocol provided by the manufacturer. The cells were incubated in the presence of EDU for 18 hours and then the samples were washed with DPBS without Ca^2+^ and Mg^2+^, followed by fixation with 3.7% formaldehyde for 15 min at room temperature. After permeabilization using 0.5% Triton X-100, the samples were incubated at room temperature for 20 min and washed with 3% bovine serum albumin (BSA) twice. Then the Click-iT reaction cocktail was added, followed by 30 min of incubation and followed again by several washes with 3% BSA. An additional stage was performed for nucleus staining with Hoechst 33342 for 30 min, followed by extensive washing with DPBS. A Nikon Eclipse Ti2 microscope at 10X was used to take photographs of the cells. A filter with excitation/emission of 495/519 nm was used to detect Alexa Fluor 488. Images were analyzed using ImageJ software to determine the percentage of EdU positive nuclei as a measure of proliferation.

### Cell morphology

hMSCs (3,000 cells/cm^2^) were seeded onto each hydrogel surface and untreated wells of a Nunclon Delta Surface clear 96 well-plate (TCP). A peptide-crosslinked hydrogel and TCP were used as controls. All the conditions were assessed in the presence and absence of IFN-γ at a concentration of 50 ng/mL. After 1 day of culture, the medium was removed, and the cells were fixed with a 4% paraformaldehyde solution for 15 minutes. Next, the samples were washed several times with DPBS before permeabilization using 0.2% Triton X-100 solution (Fisher Scientific, cat. #BP151-500) for 10 minutes. Then, hMSCs were stained with ActinRed 555 (Invitrogen™, Cat. # R415, 1:100 dilution) by incubating for 30 minutes followed by incubating with DAPI (Roche, Cat. # 10236276001, 1:1000 dilution) for 10 minutes. All conditions were washed thoroughly with DPBS before and after incubating with both dyes. Fluorescent images were obtained using a Nikon Eclipse Ti2 microscope at 20X. Measurements of the cell area and circularity in the different conditions were made using ImageJ software. Over 45 cells were counted for each condition across 4 hydrogel replicates.

### IDO activity assay

hMSCs (3,000 cells/cm^2^) with and without IFN-γ supplementation were seeded on each surface prepared in a 96 well-plate. TCP and peptide-crosslinked hydrogels were used as controls. All the conditions were assessed in the presence and absence of IFN-γ at a concentration of 50 ng/mL. IDO (Indoleamine 2,3-dioxygenase) activity was measured using a protocol previously described.^[40–42]^ Briefly, 100 μL of media were collected from each well after 3 days of culture in the specified conditions and mixed with 100 μL standard assay mixture consisting of potassium phosphate buffer (50 mM, pH 6.5), ascorbic acid (40 mM, neutralized with NaOH, Sigma, Cat. #50-81-7, Molecular Weight: 176.12), catalase (200 μg/ml, Sigma, Cat. #9001-05-2), methylene blue (20 μM, Sigma, Cat. #:122965-43-9, Molecular Weight: 319.85), and L-tryptophan (400 μM, Sigma, Cat. #73-22-3, Molecular Weight: 204.23). To allow IDO to convert L-tryptophan to N-formyl-kynurenine, the mixture was incubated at culture conditions (37°C and 5% CO_2_, protected from light) for 30 min. Next, the reaction was stopped by adding 100 µL of 30% (wt/vol) trichloroacetic acid (Sigma, Cat. #76-03-9, Molecular Weight: 163.39) and incubated at 58°C for 30 min. Solutions were then centrifuged at 10,000 RPM for 10 min. 100 μL of the supernatant was collected and mixed with 100 μL of Ehrlich’s reagent (2% (w/v) p-dimethylaminobenzaldehyde in acetic acid, (Sigma, Cat. #100-10-7, Molecular Weight: 149.19).) and incubated for 10 min. Absorbance was measured using a Synergy H1 Hybrid Multi-Mode Reader at a wavelength of 490 nm. To convert from absorbance to concentration of N-formyl-kynurenine, a standard curve varying the concentration of kynurenine (Sigma, Cas No: 2922-83-0) was also measured after undergoing the same procedure (**Figure S5**).

### Statistics

The results are presented as a mean ± standard deviation. Sample size for each experimental group is reported in the appropriate figure legend. Comparisons among groups were performed by one-way analysis of variance (ANOVA) with post-hoc Tukey HSD test. A p-value <0.05 was considered statistically significant.

## Results and Discussion

### Design of Peptoid Crosslinkers with Distinct Chain Structures

Peptoid length and secondary structure were both varied to investigate their impact on the resulting hydrogel mechanics. With respect to secondary structure, helical, non-helical, and unstructured peptoids were synthesized (**Figure 1A**). Helicity is an interesting target structure as it has been shown that helical peptoids have greater persistence lengths than similar non-helical sequences,^[24,43]^ and we previously found a marked departure from the storage moduli predicted by rubber elasticity theory for helical peptoid crosslinked PEG hydrogels.^[31]^ Peptoid helicity is induced through the inclusion of bulky, chiral monomers of a single enantiomeric state, leading to increased *cis* amide content and decreased conformational flexibility.^[21,44]^ We hypothesized that by changing the chirality of one of the submonomers used to a racemic analog, the helical secondary structure would be disrupted, but the molecule would still contain chiral character. The unstructured peptoid, meanwhile, lacks these bulky, chiral side chains, and thus no higher order structure was expected. With respect to crosslinker length, peptoids were synthesized to contain either 8 residues or 14 residues, enabling investigation of bulk mechanical scaling for a particular structure. Thus, we refer to our helical crosslinkers as H8 or H14, non-helical crosslinkers as N8 or N14, and unstructured peptoids as U8 or U14.

**Figure 1.**
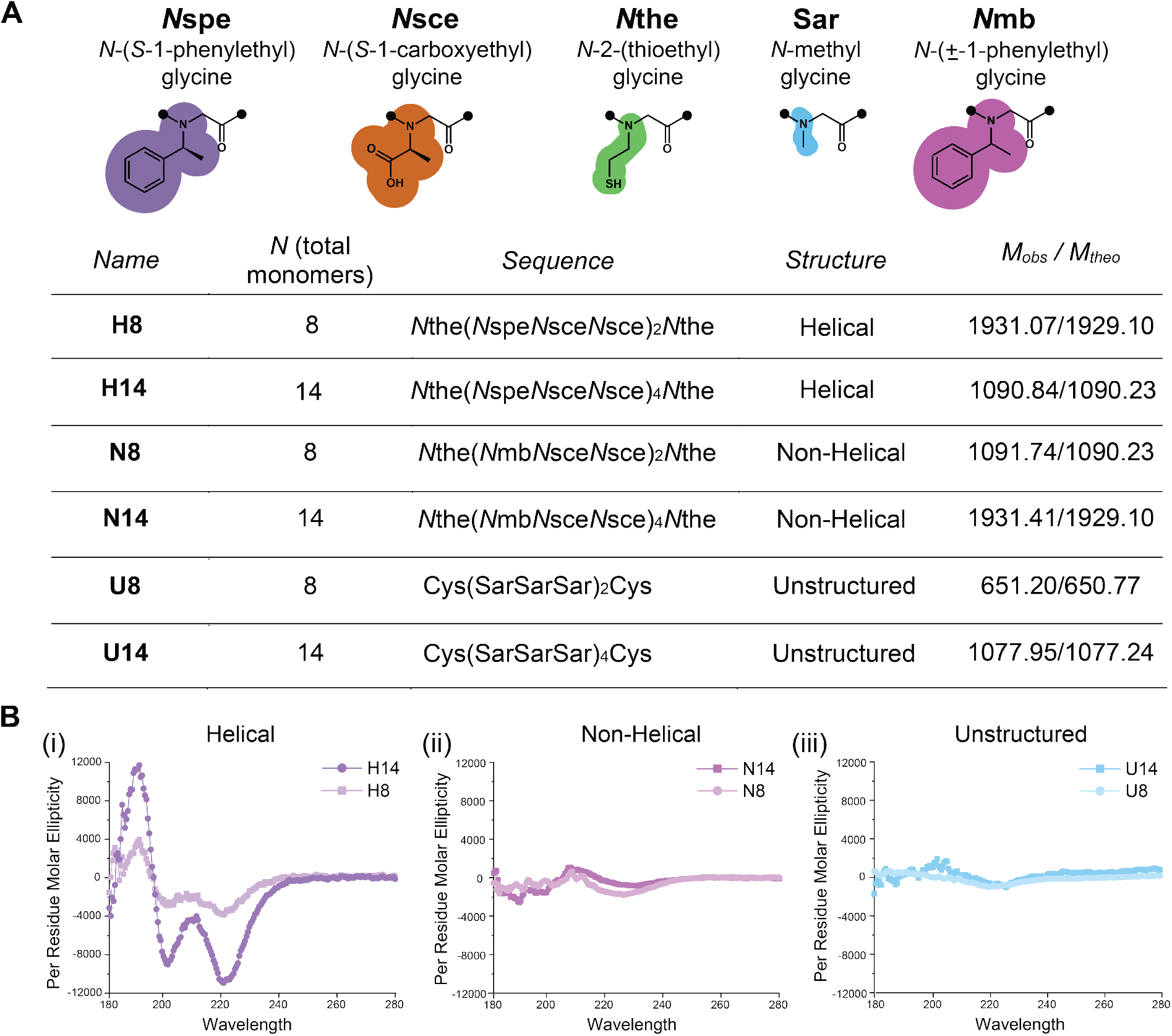
(A) N-substituted glycine residues, sequences, and masses for each peptoid crosslinker. “Cys” denotes the amino acid cysteine. (B) Circular dichroism for each peptoid illustrating characteristic peaks for the (i) helical peptoids and no peaks for the (ii) non-helical or (iii) unstructured peptoids.

For all of the crosslinkers synthesized, thiol-containing residues were included on the termini for use in thiol-ene crosslinking in the final hydrogel formulation. In addition, a peptide crosslinker (KCGGIQQWGPCK) was synthesized as a control, and a thiolated cell adhesive peptide (GCGYGRGDSPG) was synthesized for in vitro cell culture studies. The peptide crosslinker was selected with a random internal sequence to avoid secondary structure (confirmed by circular dichroism in **Figure S6**), while still possessing its inherent chirality. All peptides and peptoids were confirmed to be highly monodisperse by MALDI and analytical HPLC (**Figures S1, S2, and S3**).

Next, circular dichroism was used to probe the secondary structure of each peptoid. H8 and H14 had clear helical signatures with negative peaks at 220 and 200 nm, as well as a positive peak at 190 nm (**Figure 1Bi**).^[31,45]^ These spectra resemble peptide α-helices, with the negative peak at ∼220 nm corresponding to the nπ* transition of the amide chromophore and the other peaks corresponding to the high and low wavelength components of the exciton split ππ* transition, as previously described.^[16]^ Importantly, the per residue molar ellipticity, a measure of the robustness of the helicity, increased with peptoid length, a trend that has been previously observed for peptoid helices with different side chains.^[19,21]^ As expected, neither N8 and N14 with the racemic substitution nor U8 and U14 showed absorption profiles indicative of higher order structure (**Figure 1Bii-iii**). However, it is important to note that the lack of CD signal does not definitively disprove structure formation; rather, it confirms the lack of net molecular chirality. Thus, to examine how each sequence impacts bulk mechanics, we next incorporated these crosslinkers into hydrogels.

### Crosslinker Structure Controls Bulk Hydrogel Mechanics

NorHA hydrogels were chosen because hyaluronic acid (HA) is a well-conserved essential component in the ECM, where it is involved in cellular signaling, wound repair, and matrix organization.^[46,47]^ HA has become increasingly popular as a biomaterial building block in recent years due to its facile, tunable functionalization,^[46]^ including with norbornene groups.^[48]^ Additionally, it is degraded naturally *in vivo* by hyaluronidase, making this system particularly versatile in its applications.^[49,50]^ Peptoids have previously been incorporated into hydrogels using small molecule gelators or physical interactions, allowing the formation of soft materials.^[51–53]^ Here, we leveraged chemical crosslinking of short peptoids in a macromer system to expand the range of achievable moduli while preserving the sequence definition of the peptoid sequences.

Prior to hydrogel fabrication, we assessed the functionality of our material precursors. Norbornene-functionalized hyaluronic acid (NorHA) was synthesized to an approximate functionalization of 32% as confirmed by ^1^H NMR (**Figure S4**). For the peptoid crosslinkers, Ellman’s test was conducted for each sequence to measure the availability of free thiols and indicated a range of thiol availability from 57-85%, depending on the peptoid investigated, with some batch to batch variability (**Table S1**). MALDI spectra (**Figure S1**) indicated pure product and no presence of incomplete thiol deprotection or disulfide formation for any of the peptoids. Notably, the helical and non-helical peptoids all showed approximately 60% thiol availability, while the unstructured peptoids had higher thiol availability. We thus hypothesize that this discrepancy may due to bound water confounding the weighed mass of the peptoids, with the carboxylic acid groups on the helical and non-helical sequences leading to higher water content. The reduced thiol content was accounted for by increasing the concentration of the crosslinker until 1:1 stoichiometry was achieved for each formulation. For example, the H14 peptoid was calculated to contain 69% of the thiol necessary for full crosslinking, so a correction factor of 1.45 (100%/69%) was incorporated in the calculations, resulting in 45% more crosslinker being added to ensure proper stoichiometry and ensure equivalent crosslinking for each hydrogel.

To formulate the hydrogels, NorHA and the peptoid or peptide crosslinker were dissolved in a DMSO/water (50:50) mixture at the desired concentration with LAP photoinitiator and reacted under 365 nm light (10 mW/cm^2^, 3 min.) (**Figure 2A**). Photoinitiated thiol-ene chemistry was selected for its fast kinetics, spatiotemporal control, and ease of incorporation.^[54,55]^ All hydrogel formulations reacted quickly, reaching >95% of the final plateau modulus in seconds of photoinitiation (**Figures 2B and S7**). H14, N14, and U14 all reached 95% of their final plateau in <10 seconds, indicating no significant differences in reaction kinetics between the different secondary structures. N8 and U8, however, showed slower gelation kinetics and some physical association with the HA macromer prior to photoinitiation. Thus, we note the 8-mer peptoids may show reaction inefficiencies in crosslinking with HA and therefore focused on the 14-mer peptoids for in vitro studies, as discussed below.

**Figure 2.**
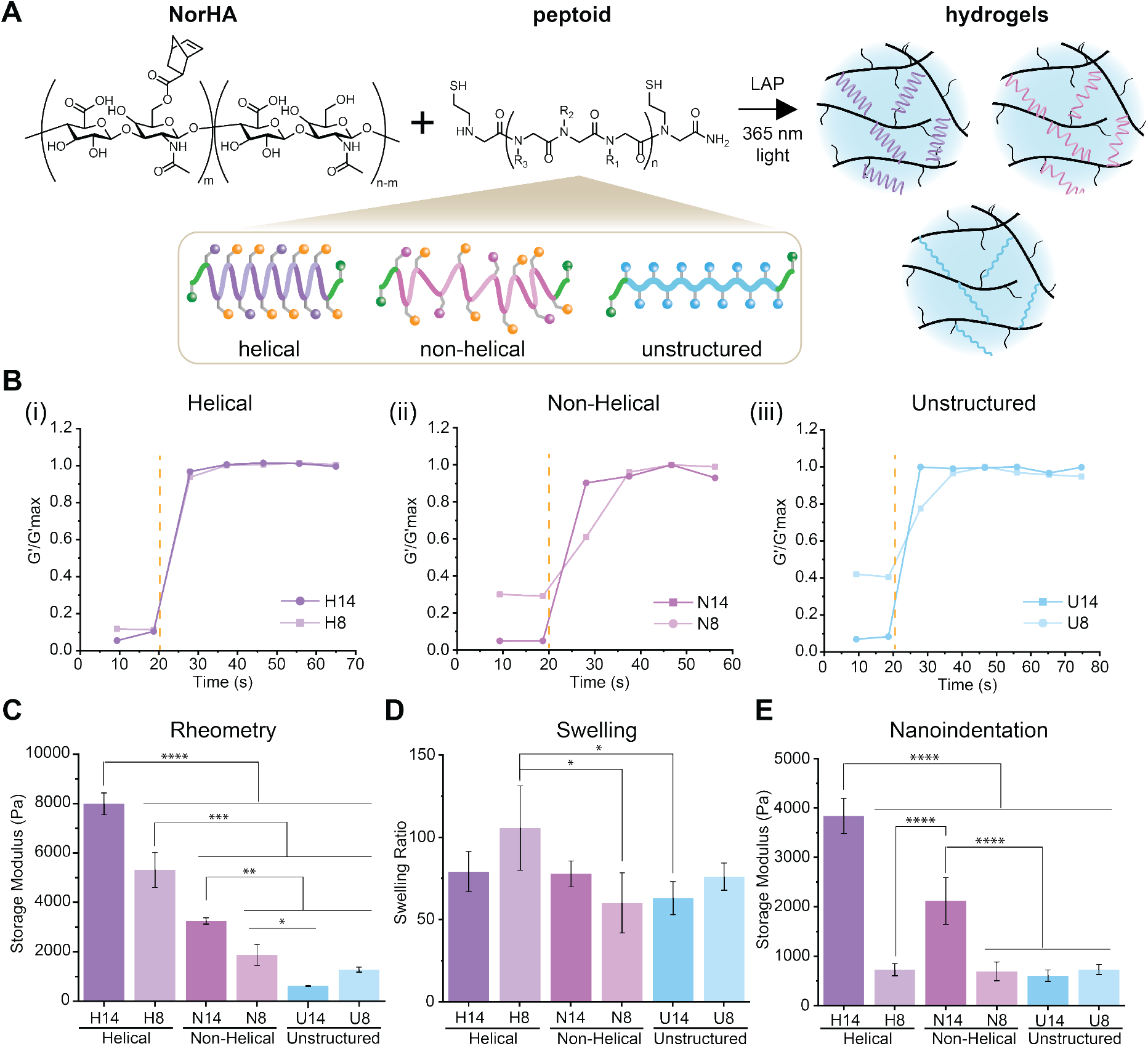
Varying peptoid secondary structure results in a range of different stiffnesses in the resulting hydrogel system, while maintaining similar swelling properties and gelation kinetics. A) Schematic representation of the photoinitiated thiol–ene crosslinking reaction between norbornene-modified HA macromers and helical peptoid crosslinkers to form a hydrogel network. B) Normalized time sweeps for each peptoid crosslinked hydrogel via shear rheometry illustrating similar kinetics. Light initiation is indicated with the orange dashed line. C) Storage moduli collected via oscillatory rheometry for each of the 6 peptoid crosslinked hydrogels. D) Swelling ratios for each hydrogel. E) Swollen storage moduli via nanoindentation. All data presented are means ± standard deviations of n = 3 samples. * denotes p<0.05, ** p<0.01, *** p<0.001, **** p<0.0001. All statistics were calculated by one-way ANOVA with post-hoc Tukey HSD test.

The secondary structure of the peptoid crosslinkers significantly affected the resulting bulk hydrogel elasticity. Using shear oscillatory rheometry, the relaxed state storage moduli 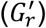 of the helical peptoids (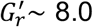 kPa for H14 and 5.3 kPa for H8) were found to be significantly greater than those of the non-helical variants (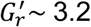 kPa for N14 and 1.9 kPa for N8) of the same length (**Figure 2C**). The unstructured peptoids yielded the softest hydrogels of all (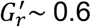 kPa for U14 and 1.3 kPa for U8). The helical peptoids likely increased the storage modulus in two ways: 1) via molecular rigidity from chirality and induced helicity, and 2) via shorter end-to-end distances due to their coiled nature. Both rubber elasticity theory and theory for networks from semiflexible polymers^[56,57]^ describe an inverse scaling relationship between the storage modulus and the mesh size (ξ), which depends on polymer end-to-end distance. Peptoid helices have been shown to be shorter than that of their non-helical counterparts; similar helices were measured to exhibit a pitch length of approximately 0.6 nm per turn, or 0.2 nm per residue.^[58,59]^ Thus, the helical peptoids are estimated to be as short as 1.6 nm to 2.8 nm for the 8-mer and 14-mer, respectively, while a fully extended peptoid would reach 2.8 nm (8-mer) or 5 nm (14-mer). Compared to other polymers, these differences are quite small,^[10]^ though would be expected to lead to increased shear modulus when comparing between different peptoid structures.

Interestingly, for the helical and non-helical crosslinkers, longer chains increased the storage moduli of the resulting hydrogels. For the helical peptoids, H14 hydrogels showed a storage modulus of 8.0 kPa compared to 5.3 kPa for H8 hydrogels (p<0.0001), and for the non-helical peptoids, N14 hydrogels showed a storage modulus of 3.2 kPa compared to 1.9 kPa for N8 hydrogels (p<0.01). This phenomenon was observed in our previous work with helical peptoid crosslinkers in PEG hydrogels, where the helical peptoid ranged from 8 to 20 monomers in length.^[31]^ This trend indicates that increased conformational stability, as shown by the larger per residue molar ellipticity in CD, yields higher molecular rigidity that dominates bulk mechanical scaling. Interestingly, although the non-helical peptoids showed no CD signature, these results indicated that their bulky, chiral side chains still confer some structure to the chains. Some of the non-helical sequences may have loose helical character due to the random incorporation of the *R* or *S* enantiomer of the *N*pe submonomer. The unstructured peptoids, however, reversed this trend and showed decreasing storage moduli with longer crosslinkers (0.6 kPa for U14 compared to 1.3 kPa for U8) though this difference was not statistically significant. Finally, as a control, a disordered peptide 12 residues in length was incorporated into HA hydrogels at the same concentration and showed a storage modulus of 2.7 kPa (**Figure S7**). Given the chiral nature of peptides, this result agreed well with the peptoid data, falling between the moduli of hydrogels with structured crosslinkers and those with achiral crosslinkers. Altogether, crosslinker structure clearly impacts bulk hydrogel mechanics in the relaxed state. Because we were interested in exploring these materials for cell culture applications, we next investigated the swelling behavior and swollen moduli for each hydrogel formulation.

### Swelling Behavior of Peptoid Crosslinked Hydrogels

The swelling ratio for each hydrogel formulation was investigated by swelling in pure water and then freezing and lyophilizing to determine the swollen and dry masses. All hydrogels swelled considerably, with each swollen hydrogel being >98% water by mass (**Figure 2D**). In addition, all hydrogels showed similar swelling ratios of 60-80, with the exception of H8. Using these swelling ratios and the measured storage moduli, 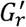, for the relaxed state, we estimated the swollen moduli for each hydrogel:

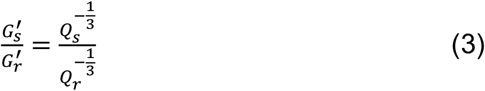

The calculated swollen moduli maintained the same trend with peptoid structure as the relaxed moduli, but decreased the differences between conditions (**Figure S8**). Importantly, the calculated swollen moduli are likely more representative of what attached cells on the hydrogels will experience *in vitro*.

To experimentally determine the swollen moduli, we used nanoindentation to measure the elastic modulus of each hydrogel in PBS (representative force curves shown in **Figure S9**). We found that H14, N14, and U14 hydrogels showed good agreement with the calculated swollen moduli, whereas the H8, N8, and U8 hydrogels did not (**Figure 2E**). In addition to the crosslinking efficiency concerns noted above, the force curves for the H8, N8, and U8 hydrogels showed significant deviations from Hertz model fits, perhaps due to the significantly softer mechanics after swelling. Thus, we chose to focus on the H14, N14, and U14 hydrogels moving forward, which presented a range of stiffness relevant for hMSC culture on hydrogel surfaces.^[33]^

### hMSC Viability and Proliferation on Peptoid-Crosslinked Hydrogels

Human mesenchymal stromal cells (hMSCs) were chosen to investigate whether the achieved range of moduli could affect cell behavior, since they have been shown to have stiffness-dependent behavior.^[33,60]^ Specifically, matrix mechanics impact the secretion of paracrine signaling factors such as indoleamine-2,3-dioxygenase (IDO), a measure of the immunomodulatory potential of hMSCs.^[32,61]^ To investigate the potential of our substrates to influence hMSC immunomodulatory behavior, we cultured cells with and without IFN-γ, since it is a cytokine known to stimulate expression of IDO (**Figure 3A**).^[35,36,61,62]^ First, metabolic activity was assessed via MTT assay after three days of culture. hMSCs were seeded on each peptoid crosslinked hydrogel with and without IFN-γ supplemented at 50 ng/mL in the cell culture medium.^[35,63]^ All hydrogels were functionalized with the cell adhesive peptide by incorporating the peptide in the macromer solution at 2 mM, which enabled simultaneous reaction via photoinitiated thiol-ene chemistry during crosslinking. The peptide crosslinked hydrogel was included as a positive control, since this formulation has been shown to be cytocompatible.^[64]^ The fluorescence intensity of the peptide hydrogel control was normalized to 100%, and all other conditions were assessed in reference to this positive control (i.e., results from peptoid-crosslinked hydrogels with IFN-γ were normalized to the results from the peptide-crosslinked hydrogel with IFN-γ) (viability raw data available in **Figure S10A**). The data from two independent trials using two separate hMSC donors indicated high metabolic activity on all hydrogel surfaces (**Figure 3B** and **Figure S10B**). The presence of IFN-γ had no significant effects on metabolic activity for any condition.

**Figure 3.**
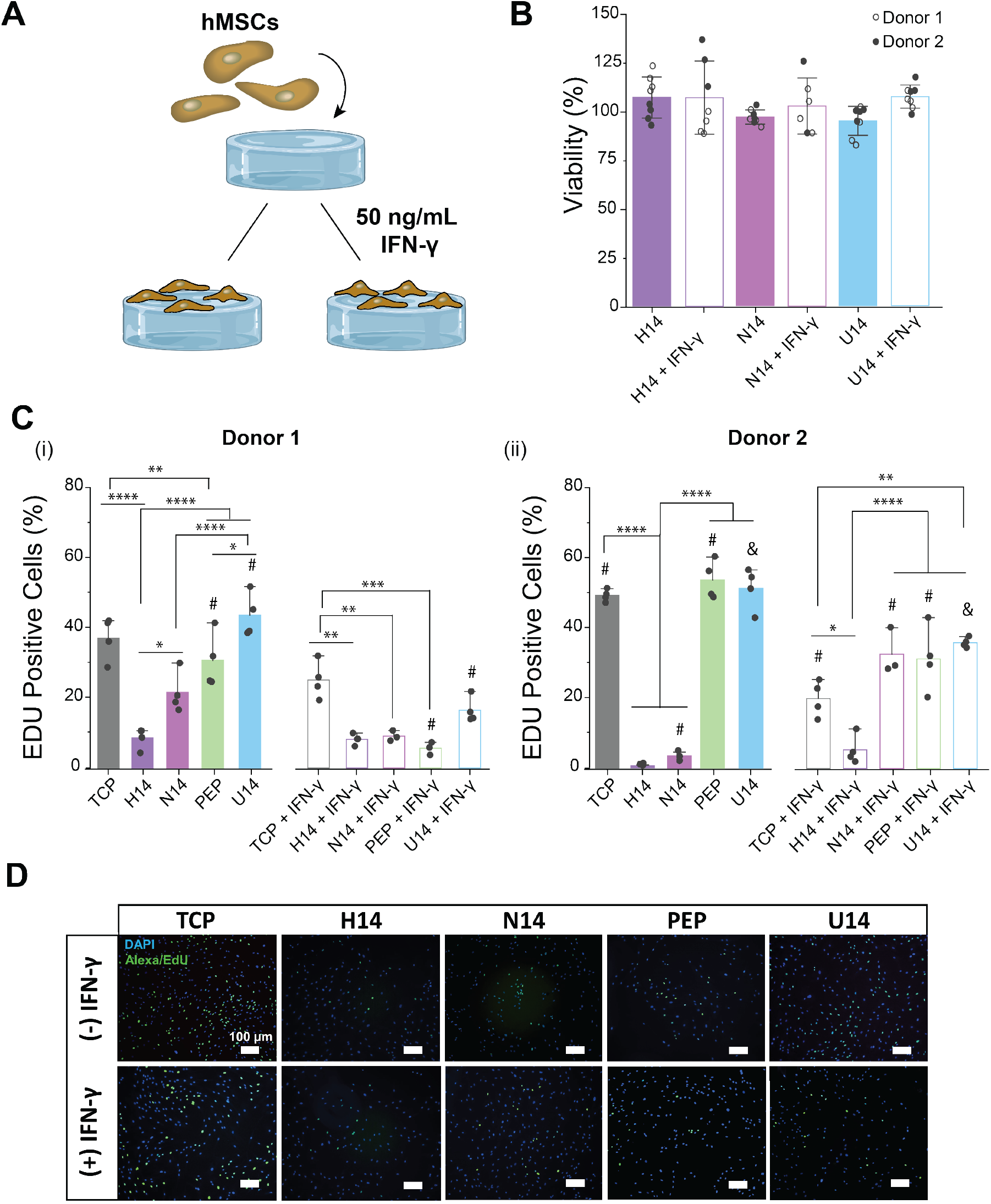
All the 14-mer peptoid crosslinked hydrogels were viable cell culture platforms for hMSC, but they resulted in significantly different levels of proliferation. A) Schematic illustrating peptoid crosslinked hydrogel formation and subsequent hMSC seeding. B) Viability of hMSCs seeded on each 14-mer peptoid crosslinked hydrogel condition after 3 days of culture with and without IFN-γ supplementation, collected via MTT assay. Unfilled circles are the data points from donor 1 and filled circles are the points from donor 2. C) Proliferation quantified by staining with EDU and counting the number of EDU positive cells after 18 hours of incubation. i) donor 1 ii) donor 2. D) Representative images from donor 1 on each condition with nuclei stained in blue and EDU positive cells stained green. The viability data is presented as means ± standard deviations of n = 4 hydrogel samples from each of two independent studies with two separate hMSC donors. All peptoid crosslinked hydrogels were normalized to the peptide crosslinked hydrogel condition. The EDU data, however, is presented as the mean ± standard deviations of n = 4 hydrogel samples with the donors separated to highlight donor to donor variability. * denotes p<0.05, ** p<0.01, *** p<0.001, **** p<0.0001. All statistics were calculated by one-way ANOVA with post-hoc Tukey HSD test. @ indicates p<0.05, & indicates p<0.01, $ indicates p<0.001, and # indicates p<0.0001 between that condition with and without IFN-γ.

To probe hMSC proliferation more directly, an EdU assay was performed on hMSCs seeded onto each hydrogel condition with and without IFN-γ supplementation after three days of culture. The percentage of EdU positive cells was determined after 18 h of incubation with the EdU solution (**Figure 3C-D**). Proliferation results demonstrated that hMSCs seeded on the hydrogel substrates show stiffness-dependent behavior. Stiffer hydrogels significantly decreased hMSC proliferation, while our softest condition (U14) presented the highest percentage of EdU positive cells (∼44% donor 1 and ∼54% donor 2 without IFN-γ) (**Figure 3C (i) and (ii)**, p<0.0001), even compared to TCP. Fewer cells were seen on the H14 and N14 conditions when imaging (**Figure 3D**). This difference was not, however, seen after seeding cells for only one day, indicating that the difference was not due to lower cell attachment to the surface, but rather a true difference in the amount of hMSC proliferation. In general, other reports have shown that hMSCs are more proliferative on stiffer hydrogels, which is opposite to the trend we observed on this system.^[32,60,65,66]^ Notably, previous work found that cellular contractility may regulate cell proliferation in an intermediate (1-5 kPa) stiffness range.^[67]^ Contractility usually increases with cell spread area on stiff hydrogels; however, it is unknown how the variation in molecular stiffness of the peptoid-crosslinked hydrogels affects contractility here.

IFN-γ supplementation has been associated with an antiproliferative effect for multiple cell types, including hMSCs.^[62,68,69]^ Our results confirmed that IFN-γ supplementation hinders hMSCs proliferation for most conditions (p<0.0001 for TCP donor 2, peptide donor 1 and 2, and U14 donor 1, p<0.01 for U14 donor 2), though a significant increase was actually found for N14 hydrogels using hMSCs from donor 2 (**Figure 3C (i) and (ii)**, p<0.0001). Altogether, these results demonstrated that peptoid crosslinked hydrogels are viable cell culture platforms and promote hMSC proliferation at softer moduli.

### Cell Morphology on Peptoid-Crosslinked Hydrogels

Next, we investigated whether the achieved range in mechanics affected attached cell morphology. Substrate stiffness has been demonstrated to play an important role in cellular mechanotransduction, in which cells transform mechanical information into biochemical signaling, thereby eliciting specific cellular behaviors such as spreading, migration, and stem cell differentiation.^[70,71]^ Several groups have reported that stiff hydrogels promote focal adhesion growth and elongation of hMSCs in particular, independent of hydrogel composition.^[33,72]^ Typically, hMSCs respond to substrate elasticities in the range of 1 kPa – 100 kPa, which overlaps with our achieved range of moduli. Thus, the hMSCs were cultured for one day on each peptoid crosslinked hydrogel formulation, then stained and imaged to measure their cytoskeletal area and circularity. TCP and the aforementioned peptide crosslinked hydrogel were used as controls. All of the conditions were assessed with and without IFN-γ supplemented in the cell culture medium. Two independent studies were conducted utilizing both hMSC donors (**Figure 4**).

**Figure 4.**
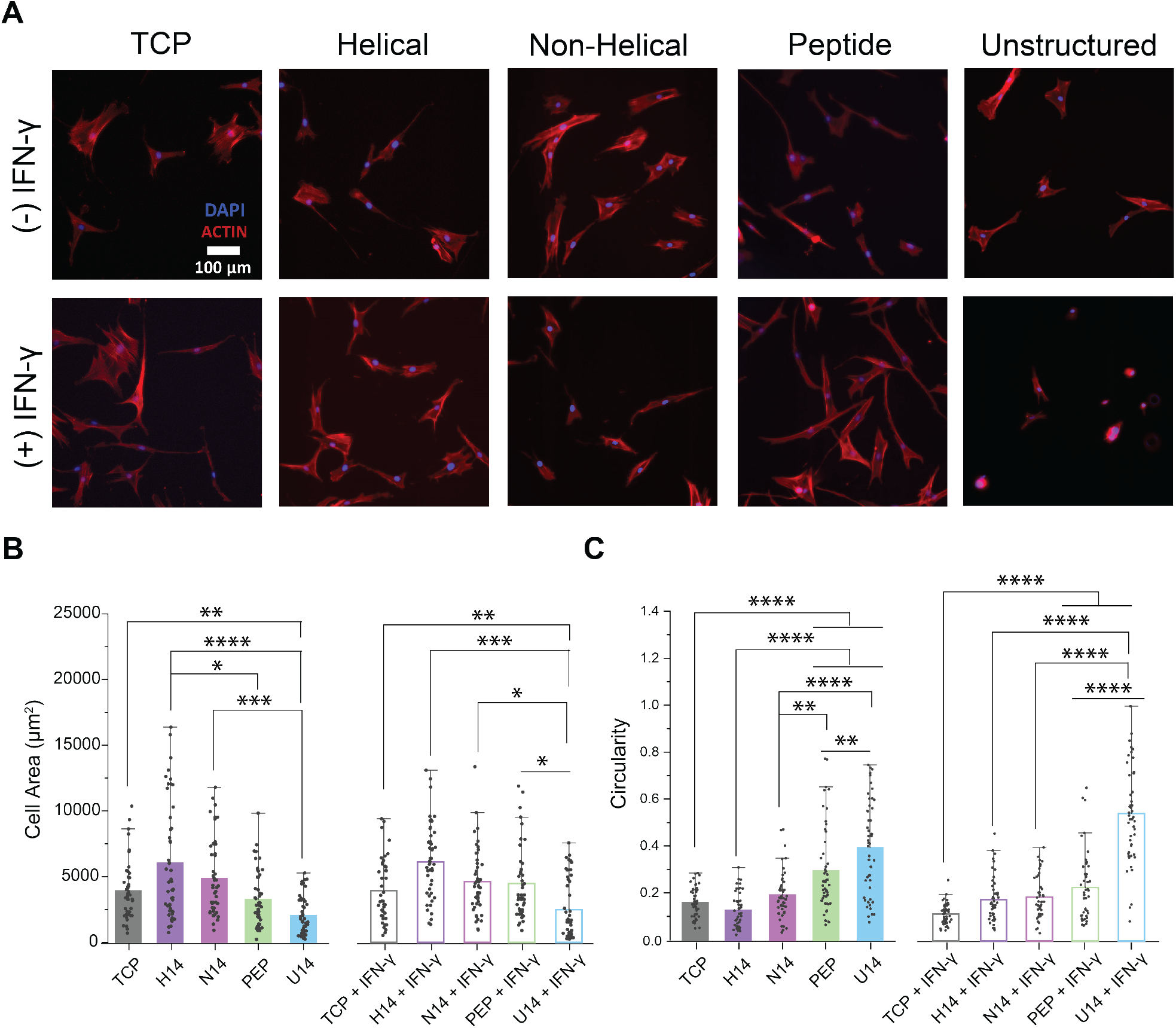
Morphological features of hMSCs cultured on TCP, peptide crosslinked hydrogels, and each 14-mer condition after one day of culture with and without IFN-γ supplementation. A) Representative fluorescent images of the hMSCs with F-actin stained with rhodamine-phalloidin (red) and nuclei stained with DAPI (blue). Scale bar 100 μm. B) Measurements of hMSC cytoskeletal area on each substrate. C) Measurements of hMSC circularity on each hydrogel substrate. All data presented are means ± standard deviations of n >45 cell measurements of two independent studies from two hMSCs donors across 4 hydrogel replicates. * denotes p<0.05, ** p<0.01, *** p<0.001, **** p<0.0001. All statistics were calculated by one-way ANOVA with post-hoc Tukey HSD test.

A clear trend was present indicating that surfaces with higher modulus increased cell area (**Figure 4A-B**), with TCP, H14, and N14 significantly increasing cell area in comparison to the U14 (p<0.01, p<0.0001, p<0.05, respectively) crosslinked hydrogels without IFN-γ. The H14 hydrogels also had significantly larger cells than the peptide crosslinked hydrogels (p<0.05). In the presence of IFN-γ, only the U14 hydrogels had significantly smaller areas than all other conditions (p<0.01 for TCP, p<0.001 for H14, p<0.05 for N14 and peptide crosslinked hydrogels). Unsurprisingly, there were no significant differences between the H8, N8, and U8 hydrogels since they all had similar swollen storage moduli (**Figure S11**). We note, however, that for a given secondary structure (e.g., H14 vs H8), the stiffer hydrogels (H14, N14, U8) resulted in more spread cells. Lastly, there were no significant differences with and without IFN-γ for any condition.

Additionally, softer hydrogels increased hMSC circularity (**Figure 4C**), both in the presence and absence of IFN-γ supplementation. Without IFN-γ supplementation, the U14 hydrogels significantly increased circularity in comparison to TCP (p<0.0001), H14 (p<0.0001), N14 (p<0.0001), and peptide (p<0.01) conditions. The peptide crosslinked hydrogel also significantly increased circularity compared to TCP (p<0.0001), H14 (p<0.0001), and N14 (p<0.01). With IFN-γ supplementation, circularity on U14 hydrogels was higher than all other conditions (p<0.0001) and the peptide crosslinked hydrogels yielded cells that were significantly more circular than TCP (p<0.0001). When comparing conditions with and without IFN-γ supplementation, there was no significant difference in circularity for any of the conditions. Previous studies have reported hMSC morphological differences in the presence of IFN-γ,^[73]^ however these changes were not significant enough to be reflected in our quantification. Altogether, our findings demonstrate that hMSC morphology can be modulated by substrate stiffness, and, importantly, that these effects are not coupled to a change in network connectivity in the hydrogel.

### Immunomodulatory Potential of hMSCs Cultured on Peptoid-Crosslinked Hydrogels

Finally, hMSCs were cultured on each substrate to determine the expression of IDO with and without IFN-γ supplementation after 3 days. IDO is an enzyme that catalyzes the degradation of L-tryptophan to produce kynurenine (by cleaving the aromatic indole ring of tryptophan), which is known to exert vital immuno-regulatory functions.^[74,75]^ IDO expression was determined by measuring the amount of N-formylkynurenine in the culture supernatant. All data were normalized by the number of cells determined by MTT assay after three days of culture to compare the relative immunosuppressive capacity of cells grown on each substrate.

In general, all hydrogels increased IDO secretion compared with culture on TCP (**Figures 5, S12**) (p<0.0001 for donor 1 without IFN-γ, p<0.001 for donor 1 with IFN-γ, p<0.001 for donor 2 with and without IFN-γ). Additionally, we found that the mechanical properties of the hydrogels may impact IDO secretion. Without IFN-γ supplementation, the softest condition (U14 hydrogels) upregulated IDO production (**Figure 5A**) (p<0.0001 compared to all other conditions), although this effect was not present in donor 2. In line with these results, previous work highlighted the effects of polyacrylamide substrate stiffness (ranging from 0.5 kPa to 200 kPa) on paracrine secretion from hMSCs, reporting that mRNA levels of cyclooxygenase-2 (COX-2), tumor necrosis factor-stimulated gene-6 (TSG-6), human leukocyte antigen-G molecules (HLA-G5), and IDO were upregulated on softer substrates after two days of culture.^[32]^ Similarly, it was reported that PEG hydrogels at different polymer concentrations (ranging from 4 to 10 wt%) with encapsulated hMSCs reported a significant decrease in IDO expression in the stiffest (10 wt%) hydrogels.^[61]^

**Figure 5.**
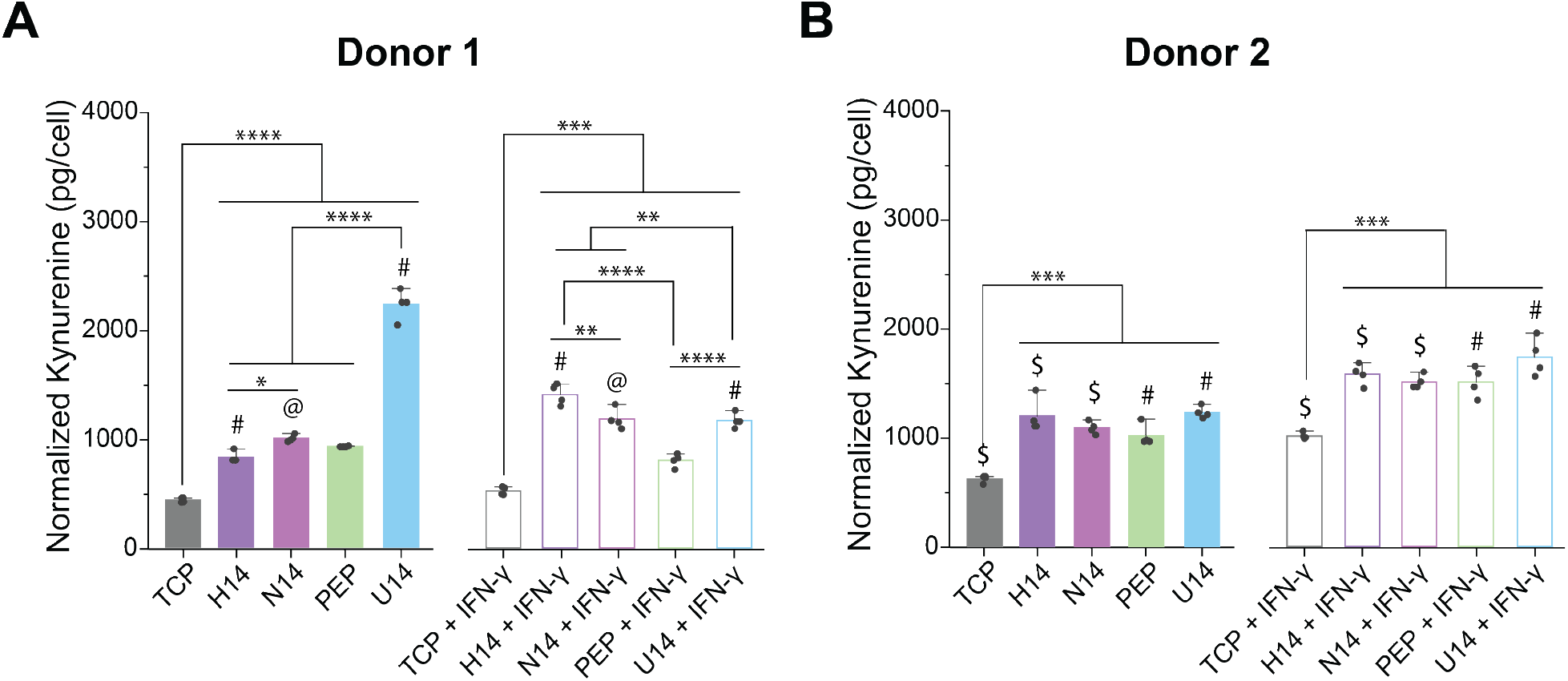
Hydrogels improve immunosuppression. The immunomodulatory potential of hMSCs cultured on TCP, peptide crosslinked hydrogels, and all the 14-mer peptoid crosslinked hydrogel conditions investigated by IDO activity with two separate hMSC donors as a measure of picograms of N-Formylkynurenine (NFK) produced with and without IFN-γ supplementation. A) Donor 1, B) Donor 2. All data presented are means ± standard deviations of n = 4 samples. * denotes p<0.05, ** p<0.01, *** p<0.001, **** p<0.0001. All statistics were calculated by one-way ANOVA with post-hoc Tukey HSD test. @ indicates p<0.05, & indicates p<0.01, $ indicates p<0.001, and # indicates p<0.0001 between that condition with and without IFN-γ.

Next, we looked at the effects of IFN-γ supplementation on IDO secretion for each hydrogel condition. For donor 1, H14 and N14 crosslinked hydrogels had significantly higher IDO production (p<0.0001 and p<0.05 respectively), but peptide crosslinked hydrogels showed no significant difference and the U14 crosslinked hydrogels actually showed a significant decrease in IDO (p<0.0001) (**Figure 5A**). For donor 2, however, the trend was much clearer. IFN-γ significantly increased IDO for each condition (p<0.001 for TCP, H14, and N14 and p<0.0001 for peptide and U14) (**Figure 5B**). The H8, N8, and U8 hydrogels, which all had similar soft swollen moduli, demonstrated no clear trend in IDO production with IFN-γ supplementation (**Figure S12**).

Altogether, these data indicate that softer hydrogel mechanics may increase IDO production up to a certain point, at which point the effect of substrate mechanics plateaus, and that IFN-γ supplementation increases IDO production on stiffer hydrogels but may have less effect on softer hydrogels. To investigate this result further, we fabricated hydrogels of varying moduli by changing the concentration of NorHA, rather than crosslinker structure (**Figure S13A**), allowing investigation of substrate moduli over an expanded range, from 200 Pa – 30,000 Pa (relaxed state storage moduli). Interestingly, the softer hydrogels again upregulated IDO production (**Figure S13B**), but the effect plateaued for the 2 wt% and 4 wt% hydrogels, which overlap in the stiffness range achieved for our peptoid-crosslinked hydrogels. IFN-γ supplementation appeared to increase IDO production over the entire stiffness range using cells from Donor 1. In previous work, IFN-γ supplementation was found to increase IDO production on soft (∼50-200 Pa) fibrin and collagen gels.^[76]^ Overall, these data merit further investigation and show that hydrogel mechanics play an important role in the resulting immunomodulatory activity of hMSCs. In addition, these data suggest that soft peptoid-crosslinked hydrogels may regulate immunomodulatory activity in a different fashion than other soft hydrogel substrates.

## Conclusion

In conclusion, we modulated the mechanical properties of naturally-derived hyaluronic acid hydrogels by crosslinking with peptoids of different secondary structure and chain stiffness. We synthesized a helical, non-helical, and unstructured peptoid sequence at lengths of 8 and 14 monomers, as well as a peptide crosslinker. Crosslinkers with increased chain stiffness significantly raised the bulk mechanics of the formed hydrogels. Our system offers a strategy for decoupling matrix mechanics from network connectivity, allowing for better recapitulation of the hierarchical nature of the ECM and an expanded toolbox for design of biophysically-relevant cell culture scaffolds. In this study, the range of developed hydrogel mechanics was significant for hMSC culture. All substrates resulted in highly viable hMSCs, with softer hydrogels leading to smaller, more circular cells that proliferated more and produced higher IDO concentrations. In addition, the softest peptoid-crosslinked hydrogels rendered the cells less sensitive to cytokine supplementation. In future work, we anticipate that the developed materials will shed insight to mechanoregulation of hMSC immunomodulatory potential and enable the development of improved strategies for hMSC manufacturing. In addition, future inclusion of molecular rigidity or crosslinker structure in scaffold design could lead to more biomimetic matrices for cell manufacturing applications, tissue engineering, and regenerative medicine.

## Supporting Information

MALDI spectrum for each peptoid sequence. Analytical HPLC spectrum for each peptoid sequence. MALDI spectrum for each peptide sequence. ^1^HNMR of Norbornene-functionalized hyaluronic acid (NorHA). Kynurenine standard curve for IDO calculations. Peptide circular dichroism. Table of thiol percentages for each peptoid crosslinker. Time sweep for the peptide crosslinked hydrogel. Calculated swollen storage moduli. Example calculation for hydrogel swollen moduli. Representative nanoindentation measurements for all the peptoid crosslinked hydrogel formulations. Viability raw data and normalized viability of hMSCs seeded on each 8-mers peptoid crosslinked hydrogel condition. Morphological features of the hMSCs cultured on each 8-mers peptoid crosslinked hydrogel condition after one day of culture with and without IFN-γ supplementation. The immunomodulatory potential of hMSCs cultured on all the 8-mer peptoid crosslinked hydrogel conditions investigated by IDO activity with two separate hMSC donors. Hydrogel mechanics (changing the wt% of NorHA) vs immunomodulatory potential of hMSCs.

## Acknowledgement

We thank Mariah Austin for help with figure production. We thank Francis Garcia for assisting in maintaining the hMSC cell culture. The graphical abstract and Figure 3A contain icons generated by BioRender.com. This research was supported by the Burroughs Welcome Fund (CASI-1015895, A. M. R.), the National Institutes of Health (R35GM138193, A.M.R), and the National Science Foundation (NSF MRSEC DMR-1720595 A. M. R.). In addition, L. D. M. was supported through a National Science Foundation Graduate Research Fellowship. The authors acknowledge the use of shared research facilities supported in part by the Texas Materials Institute, the Center for Dynamics and Control of Materials: an NSF MRSEC (DMR-1720595), and the NSF National Nanotechnology Coordinated Infrastructure (ECCS-1542159). We also acknowledge the use of shared facilities in the Targeted Therapeutic Drug Discovery & Development Program (CPRIT Core Facilities Support Award (grant #RP160657)) and the UT Proteomics Facility.

## Data Availability Statement

The data that support the findings of this study are available from the corresponding author upon request.

