## Supplemental Information for "Crosslinker Structure Modulates Bulk Mechanical Properties and Dictates hMSC Behavior on Hyaluronic Acid Hydrogels"

### Supporting Information

Figure S1. MALDI spectrum for each peptoid sequence. 2

Figure S2. Analytical HPLC spectrum for each peptoid sequence. 3

Figure S3. MALDI spectrum for each peptide sequence. 4

Figure S4. $^1$HNMR of Norbornene-functionalized hyaluronic acid (NorHA). 5

Figure S5. Kynurenine standard curve for IDO calculations. 6

Figure S6. Peptide circular dichroism. 7

Table S1. Table of thiol percentages for each peptoid crosslinker. 8

Figure S7. Time sweep for the peptide crosslinked hydrogel. 9

Figure S8. Calculated swollen storage moduli. 10

Figure S9. Representative nanoindentation measurements. 11

Figure S10. Viability raw data for all conditions. 12

Figure S11. Morphological features of the hMSCs cultured on each 8-mer condition. 13

Figure S12. Immunomodulatory potential of hMSCs cultured on each 8-mer condition. 14

Figure S13. Immunomodulatory potential of hMSCs on control HA hydrogels. 15

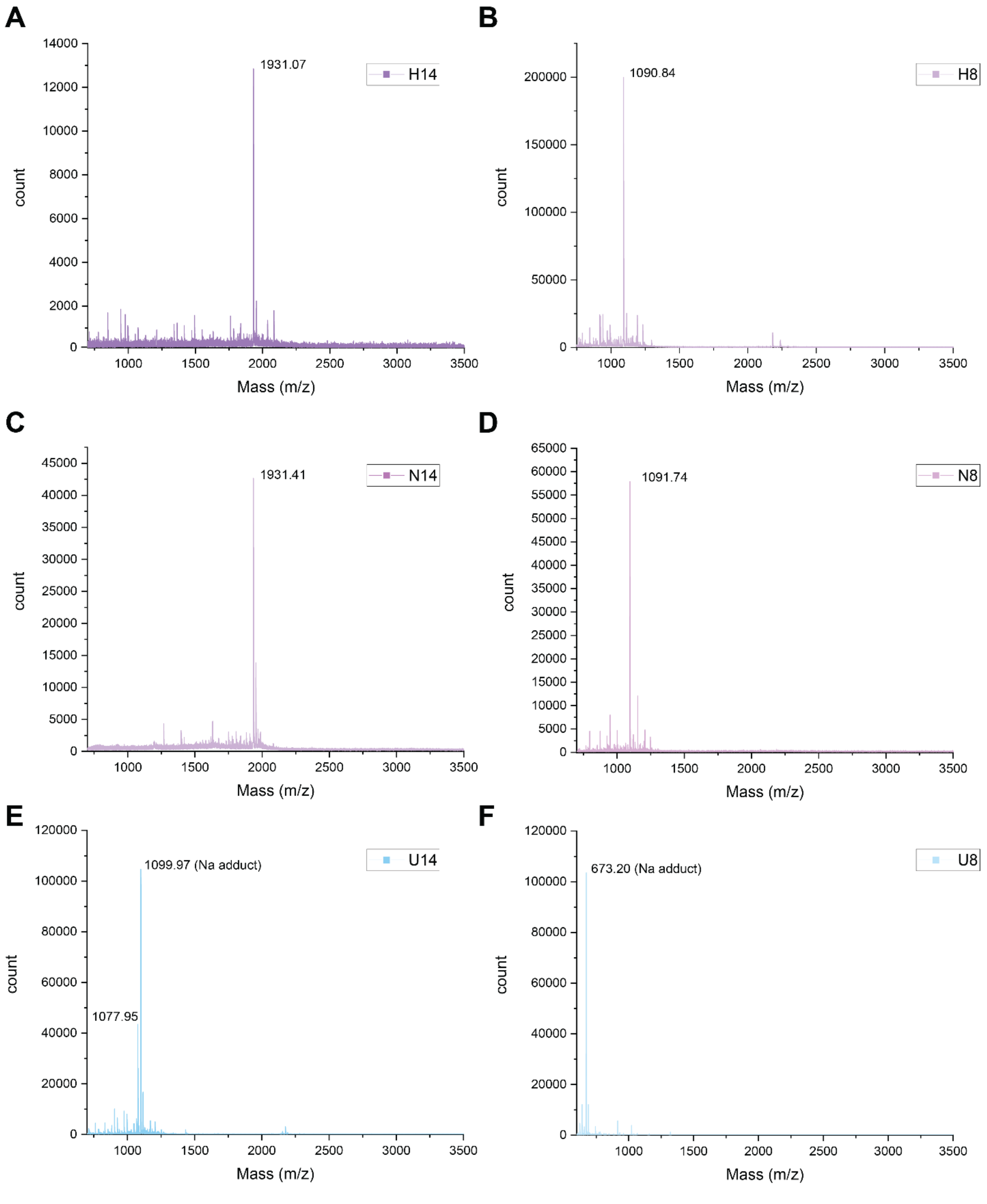


**Figure S1.** All peptoid sequences were confirmed to be highly monodisperse by MALDI. A) H14, B) H8, C) N14, D) N8, E) U14, F) U8.

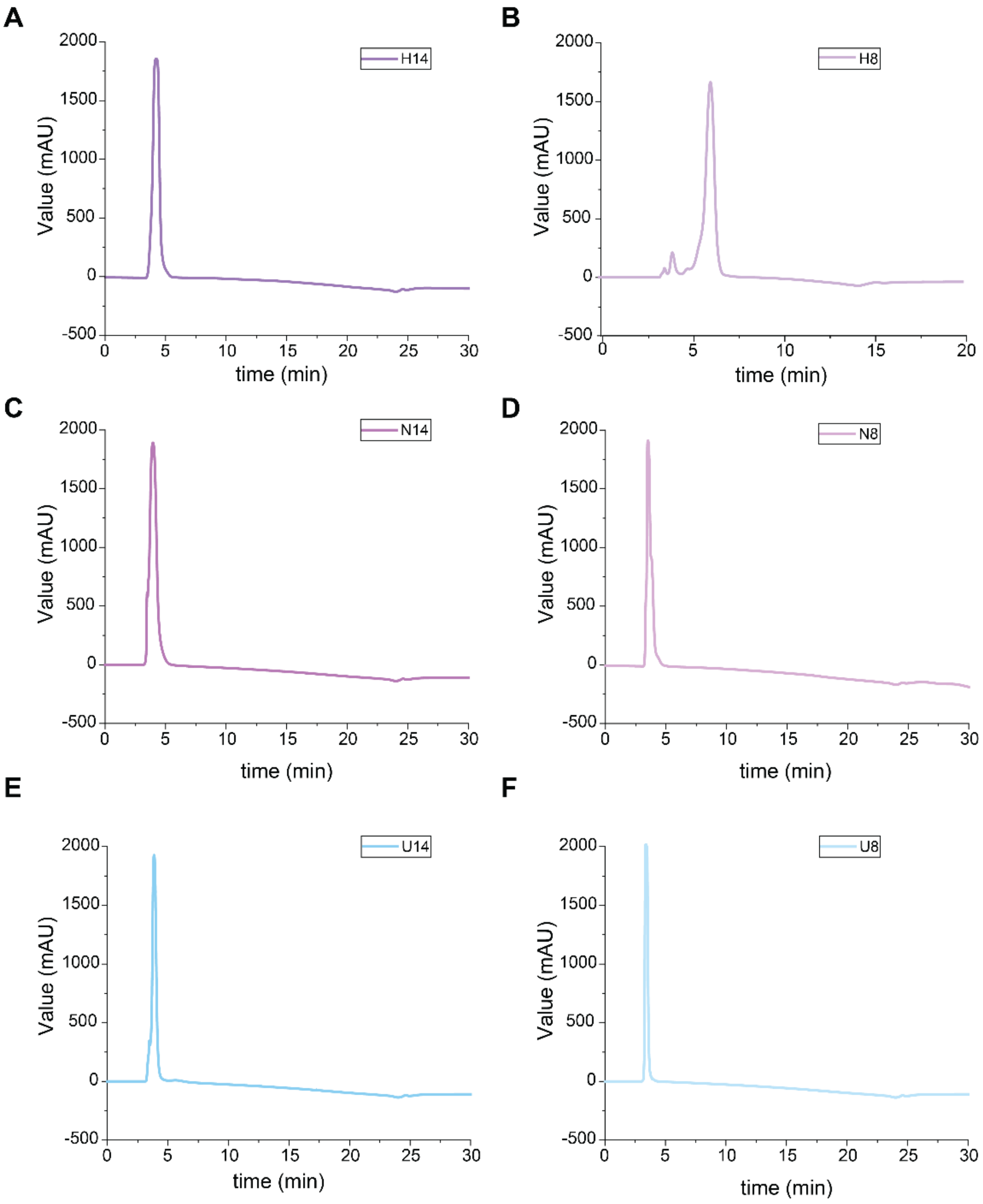


**Calculated Purity by Integration**

| Crosslinker | H14 | H8 | N14 | N8 | U14 | U8 |
|---|---|---|---|---|---|---|
| Purity (%) | 100% | 90% | 94% | 92% | 91% | 100% |

**Figure S2.** All peptoid sequences were confirmed to be highly pure by analytical HPLC. A) H14, B) H8, C) N14, D) N8, E) U14, F) U8.

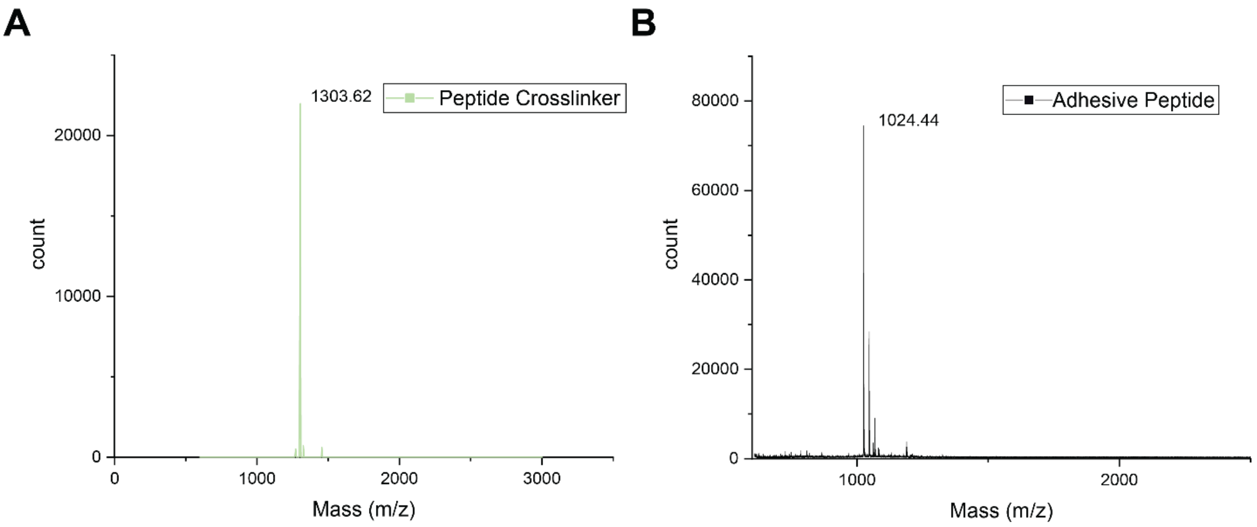


**Figure S3.** Both peptides were found to be highly monodisperse by MALDI. A) peptide crosslinker (KCGGIQQWGPCK), B) cell adhesive peptide (GCGYGRGDSPG).

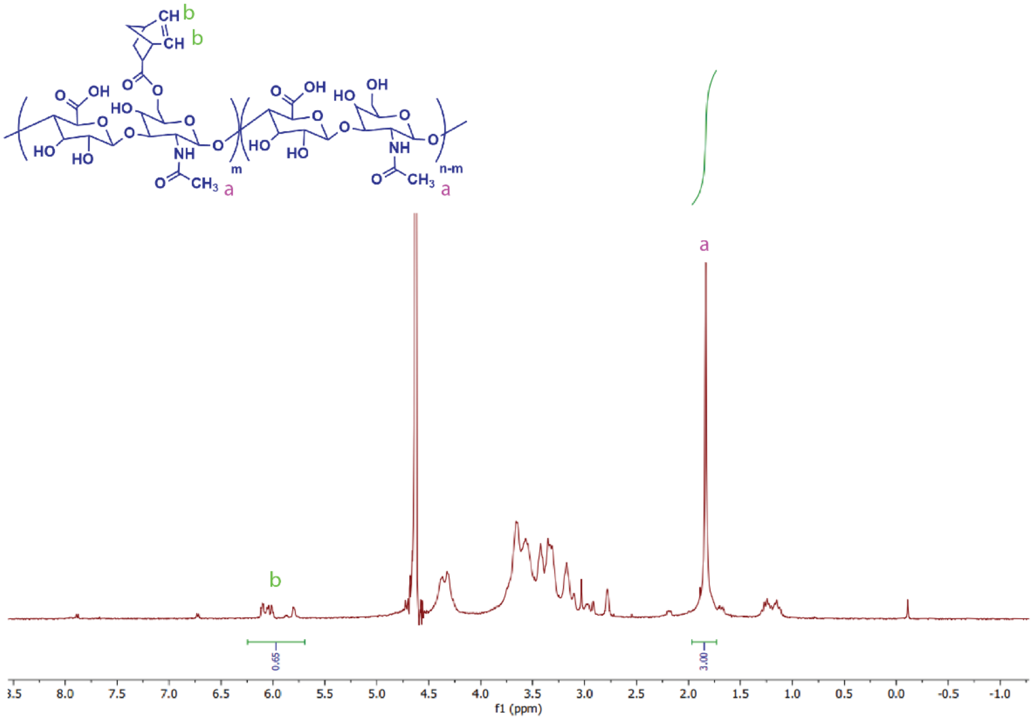


**Figure S4.** Norbornene-functionalized hyaluronic acid (NorHA) was synthesized to an approximate functionalization of 32% as confirmed by $^{1}$H NMR.

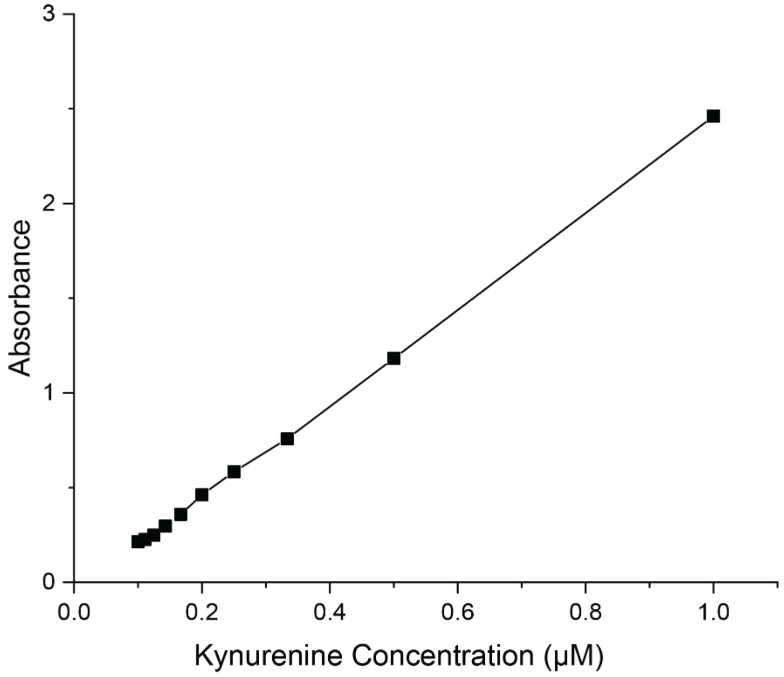


**Figure S5:** Standard curve relating kynurenine concentration to measured absorbance at 490 nm in the presence of Ehrlich's reagent.

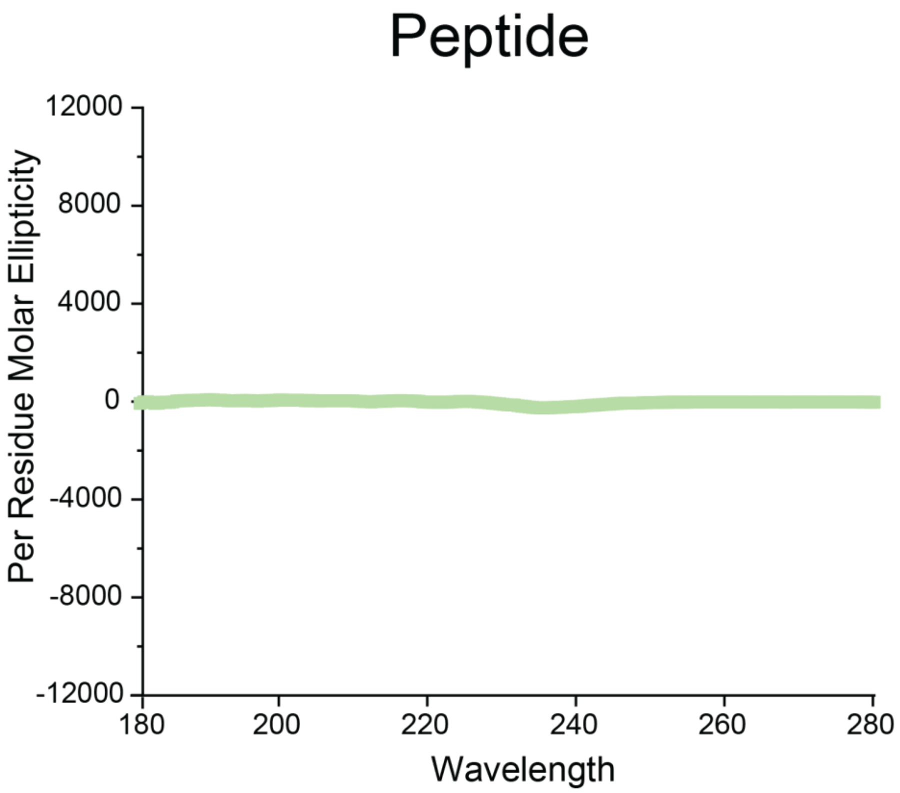


**Figure S6.** Circular dichroism of the scramble peptide, confirming no appreciable signal is present.

**Table S1.** Thiol percentages for each formulation.

| Peptoid | H14 | H8 | N14 | N8 | U14 | U8 |
|---|---|---|---|---|---|---|
| Ellman's (Thiol %) | 69% | 57% | 60% | 60% | 85% | 76% |
| St Dev | 19% | 19% | 15% | 24% | 6% | 23% |

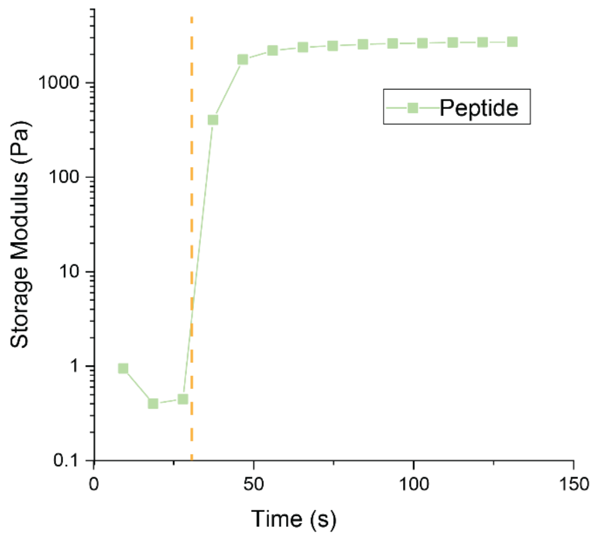


**Figure S7.** Rheometry time sweep curve for the peptide crosslinked hydrogel. Note that the UV light was not turned on until ~25s into the measurement as illustrated by the orange dashed line.

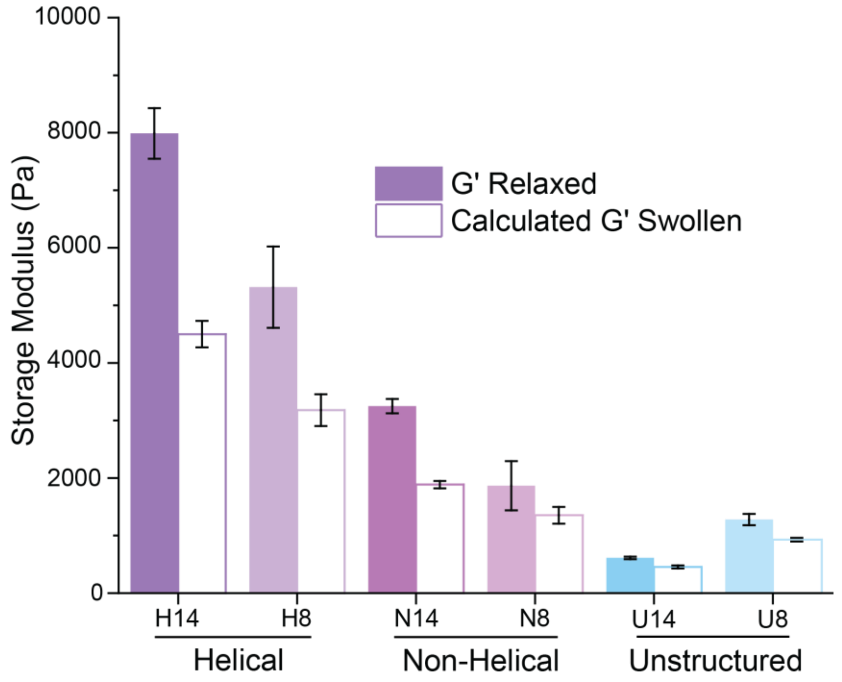


**Figure S8.** Calculated swollen storage moduli using equation 3, example calculation shown below.

**Example Calculation for determining the swollen moduli for each hydrogel using the relaxed modulus and swelling ratio.**

$$\frac{G_s}{G_r} = \frac{RT\rho_x Q_s^{-\frac{1}{3}}}{RT\rho_x Q_r^{-\frac{1}{3}}} = \frac{Q_s^{-\frac{1}{3}}}{Q_r^{-\frac{1}{3}}}$$

$$G_s = \frac{G_r Q_s^{-\frac{1}{3}}}{Q_r^{-\frac{1}{3}}}$$

$$G_{s,H14} = \frac{(7988.33\ Pa) * (79.15)^{-\frac{1}{3}}}{(14)^{-\frac{1}{3}}} = 4499\ \mathrm{Pa}$$

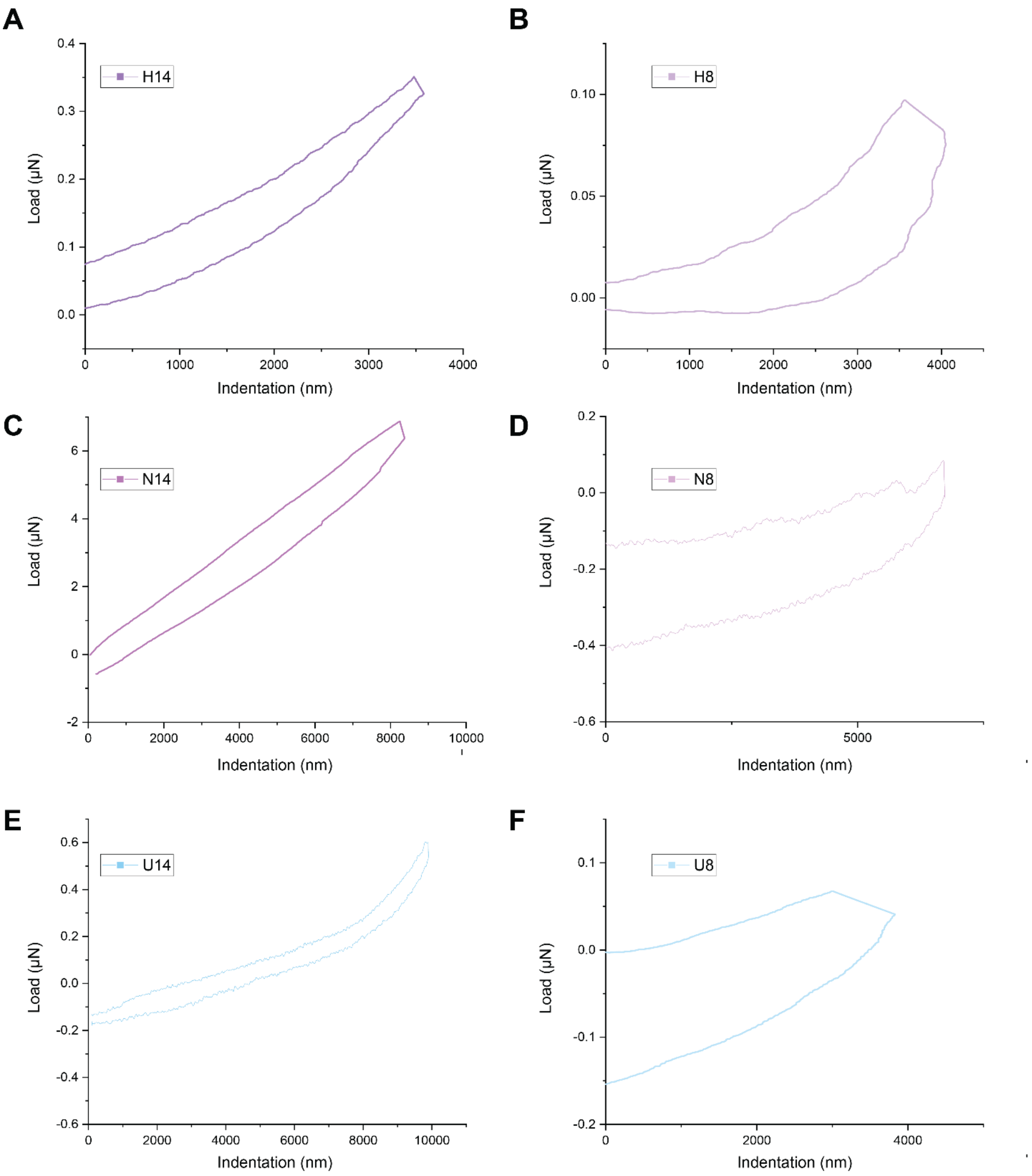


**Figure S9.** Representative nanoindentation measurements for all the peptoid crosslinked hydrogel formulations. A) H14, B) H8, C) N14, D) N8, E) U14, F) U8.

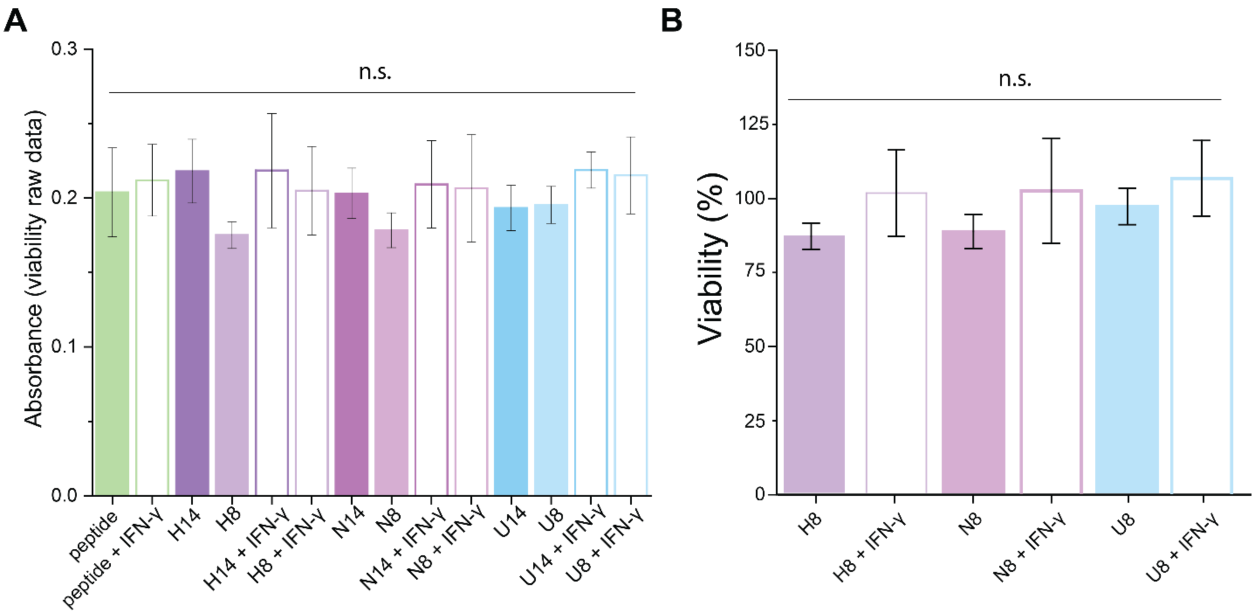


**Figure S10.** Viability of hMSCs seeded on each hydrogel condition after 3 days of culture with and without IFN-γ supplementation, collected via MTT assay. A) Viability raw data, B) Normalized viability of hMSCs seeded on each of the 8-mer crosslinked hydrogel conditions. All data presented are means ± standard deviations of n = 4 samples from each of two independent studies with two separate hMSC donors. All peptoid crosslinked hydrogels were normalized to the peptide crosslinked hydrogel condition. * denotes $p<0.05$, ** $p<0.01$, *** $p<0.001$, **** $p<0.0001$. All statistics were calculated by one-way ANOVA with post-hoc Tukey HSD test.

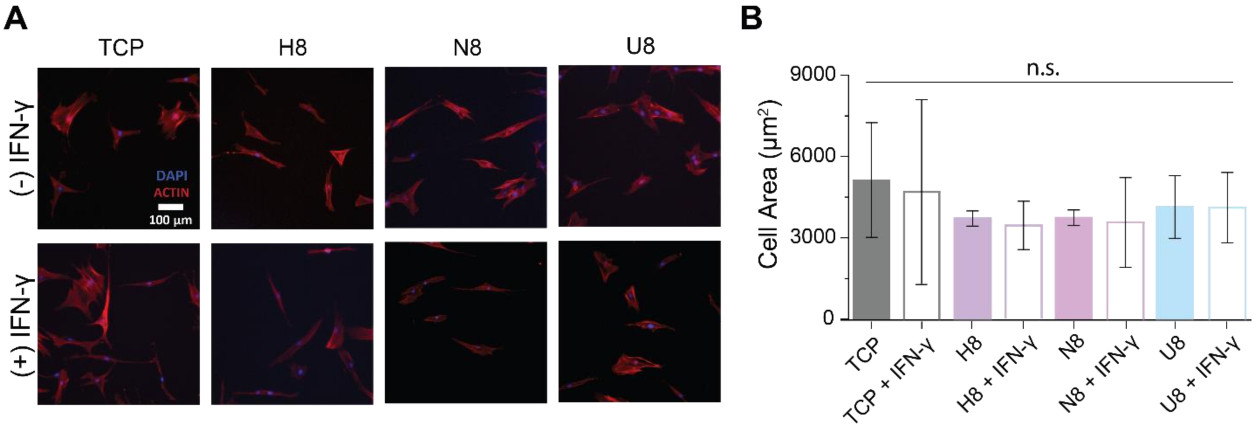


**Figure S11.** Morphological features of hMSCs cultured on each 8-mer crosslinked hydrogel condition after one day of culture with and without IFN-γ supplementation. A) Representative fluorescent images of the hMSCs with F-actin stained with rhodamine-phalloidin (red) and nuclei stained with DAPI (blue). Scale bar 100 μm. B) Measurements of average hMSC cytoskeleton surface area on hydrogels crosslinked with each peptoid 8-mer. All data presented are means ± standard deviations of n >60 measurements of two independent studies from two hMSCs donors. All data presented are means ± standard deviations of n = 4 samples. * denotes $p<0.05$, ** $p<0.01$, *** $p<0.001$, **** $p<0.0001$. All statistics were calculated by one-way ANOVA with post-hoc Tukey HSD test.

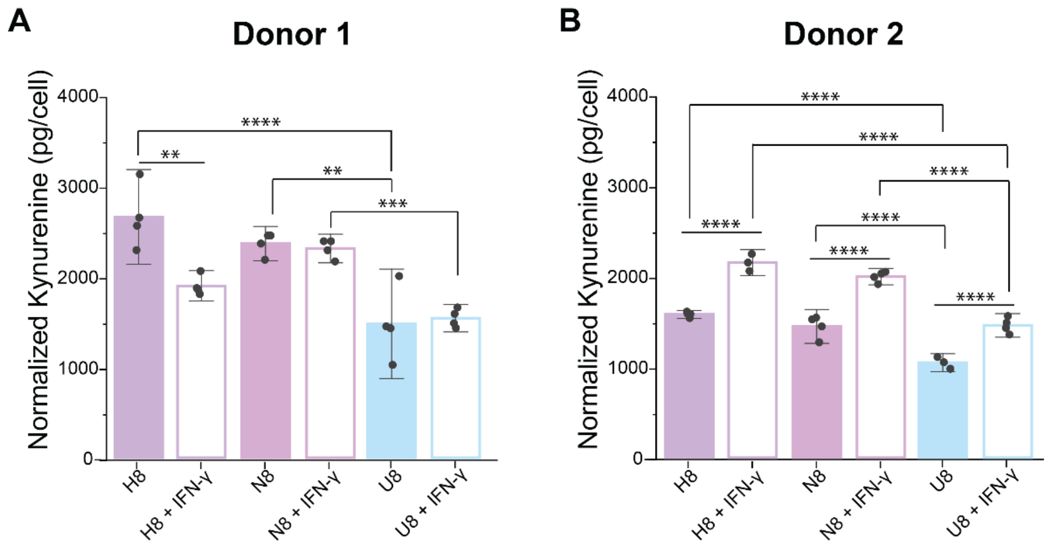


**Figure S12.** The immunomodulatory potential of hMSCs cultured on hydrogels crosslinked with each peptoid 8-mer. This was investigated by IDO activity measured in picograms of N-Formylkynurenine (NFK) produced with and without IFN-γ supplementation with two separate hMSC donors. A) Donor 1, B) Donor 2. All data presented are means ± standard deviations of n = 4 samples. * denotes $p<0.05$, ** $p<0.01$, *** $p<0.001$, **** $p<0.0001$. All statistics were calculated by one-way ANOVA with post-hoc Tukey HSD test.

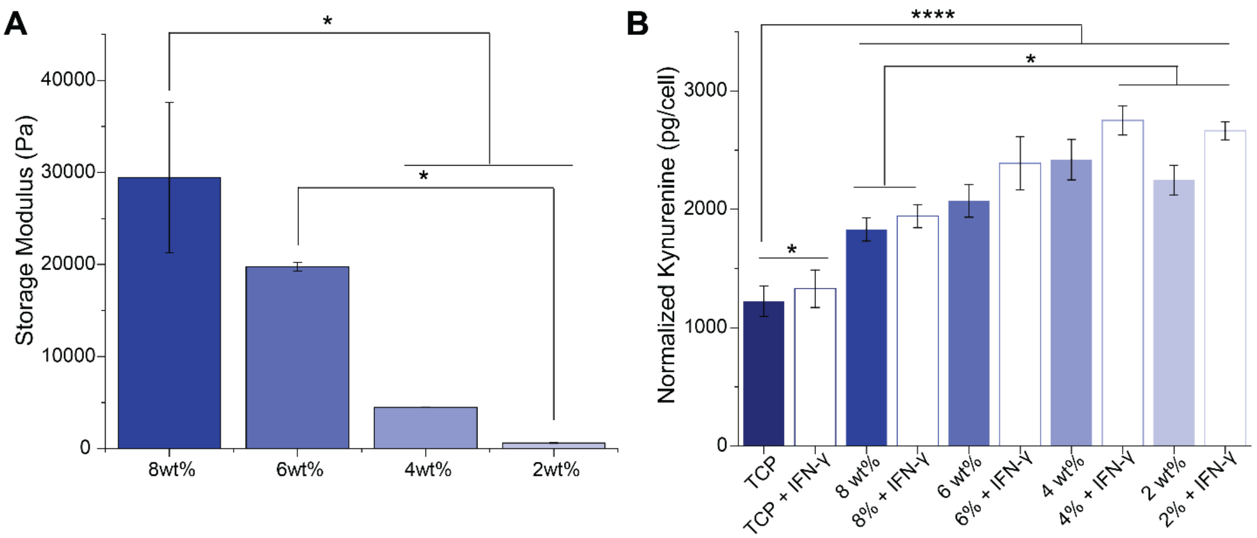


**Figure S13.** Softer hydrogels upregulate the immunomodulatory potential of hMSCs. Representative NorHA hydrogels were fabricated at different wt%'s using a simple dithiol crosslinker (1,4-Dithiothreitol). A) Storage moduli collected via shear oscillatory rheometry for different concentrations of NorHA. B) IDO activity measured in picograms of N-Formylkynurenine (NFK) produced with and without IFN-γ supplementation for Donor 1 hMSCs seeded on each surface. All data presented are means ± standard deviations of n = 4 samples for the IDO and 2 samples for the modulus. * denotes $p<0.05$, ** $p<0.01$, *** $p<0.001$, **** $p<0.0001$. All statistics were calculated by one-way ANOVA with post-hoc Tukey HSD test.
